# Competitive coordination of the dual roles of the Hedgehog co-receptor in homophilic adhesion and signal reception

**DOI:** 10.1101/2021.02.19.432013

**Authors:** Shu Yang, Ya Zhang, Chuxuan Yang, Xuefeng Wu, Sarah Maria El Oud, Rongfang Chen, Xudong Cai, Xufeng S. Wu, Ganhui Lan, Xiaoyan Zheng

## Abstract

Hedgehog (Hh) signaling patterns embryonic tissues and contributes to homeostasis in adults. In *Drosophila*, Hh transport and signaling are thought to occur along a specialized class of actin-rich filopodia, termed cytonemes. Here, we report that Interference hedgehog (Ihog) not only forms a Hh receptor complex with Patched to mediate intracellular signaling, but Ihog also engages in *trans*-homophilic binding leading to cytoneme stabilization in a manner independent of its role as the Hh receptor. Both functions of Ihog (*trans*-homophilic binding for cytoneme stabilization and Hh binding for ligand sensing) involve a region of the first fibronectin repeat of the extracellular domain. Thus, the Ihog-Ihog interaction and the Hh-Ihog interaction cannot occur simultaneously for a single Ihog molecule. By combining experimental data and mathematical modeling, we determined that Hh-Ihog heterophilic interaction dominates and Hh can disrupt and displace Ihog molecules involved in *trans*-homophilic binding. Consequently, we proposed that the weaker Ihog-Ihog *trans* interaction promotes and stabilizes direct membrane contacts along cytonemes and that, as the cytoneme encounters secreted Hh ligands, the ligands trigger release of Ihog from *trans* Ihog-Ihog complex enabling transport or internalization of the Hh ligand-Ihog-Patched -receptor complex. Thus, the seemingly incompatible functions of Ihog in homophilic adhesion and ligand binding cooperate to assist Hh transport and reception along the cytonemes.

**One-sentence summary:** Apparently incompatible functions of Ihog in homophilic adhesion and ligand binding cooperate in Hh transport and reception.

## INTRODUCTION

Hedgehog (Hh) signaling plays essential roles in patterning of multicellular embryos and maintaining adult organ homeostasis. Aberration in the precise temporal-spatial regulation and transduction of the Hh signaling pathway is involved in some birth defects (Muenke and Beachy 2000) and various proliferative disorders, such as the growth of malignant tumors (Varjosalo and Taipale 2008).

Hh protein precursor undergoes autoprocessing and lipid modification that generates the mature Hh ligand as an amino-terminal signaling peptide (HhN) dually modified by palmitoyl and cholesteryl adducts (Mann and Beachy 2004). Intracellular signaling is triggered by binding of the dually lipidated Hh ligand to the receptor. The *Drosophila* Hh receptor regulates the component, Smoothened, and limits the range of signaling by sequestering Hh ligand. The Hh receptor is comprised of Patched (Ptc) and a member of the Ihog family, which in *Drosophila* is one of the functionally interchangeable proteins encoded by *interference Hedgehog* (*ihog*) or *brother of ihog* (*boi*) (Lum et al. 2003; McLellan et al. 2006; Yao et al. 2006; Camp et al. 2010; Chou et al. 2010; Hartman et al. 2010; Yan et al. 2010; Zheng et al. 2010).

The Ihog family proteins are type I single-span transmembrane proteins with immunoglobulin (Ig) and fibronectin type III (FNIII) domains, resembling typical cell adhesion molecules in the Ig-CAM (Ig cell adhesion molecule) superfamily. Previous biochemical and structural studies showed that the first FNIII domain (Fn1) in the extracellular portion of Ihog is involved in binding to the Hh ligand (McLellan et al. 2006; Yao et al. 2006), whereas the second FNIII domain (Fn2) of Ihog interacts with Ptc. Both Fn1 and Fn2 domains are required for Hh signal reception through the formation of a high-affinity multimolecular complex of Ihog, Ptc, and Hh (Zheng et al. 2010). Ihog proteins not only play an essential role in Hh signal transduction but also mediate cell-cell interactions in a homophilic, calcium-independent manner (Zheng et al. 2010; Hsia et al. 2017; Wu et al. 2019). The region that mediates the *trans* Ihog-Ihog interaction overlaps with the region that mediates the interaction with Hh on the Ihog Fn1 domain and includes a region where the negatively charged glycan heparin binds (McLellan et al., 2006; Wu et al., 2019). Heparin is required for not only Hh-Ihog interactions but also Ihog-Ihog homophilic *trans* interactions *in vitro* (McLellan et al. 2006; Zhang et al. 2007; Wu et al. 2019).

The presence of dual functions as an adhesion protein and as a signaling protein is not unique to Ihog proteins. Other members of the Ig-CAM family, such as the netrin receptor DCC, the Slit receptor Robo, and neural cell adhesion molecule (N-CAM), have dual roles. These proteins act as molecular “glue” that holds cells together and as molecular sensors to mediate cellular responses, such as motility, proliferation, and survival (Juliano 2002; Orian-Rousseau and Ponta 2008). However, ligand binding and cell adhesion are often structurally separated and involve different extracellular domains (Frei et al. 1992; Martin-Bermudo and Brown 1999; Sjostrand et al. 2007). In contrast, the Ihog protein couples these distinct functions within the same region. The physiological consequences of coupling two distinct functions into the same region of the Ihog protein are unknown.

In the *Drosophila* wing imaginal discs, Hh is secreted in the posterior (P) compartment and spreads toward the anterior (A) compartment (Basler and Struhl 1994; Capdevila et al. 1994; Tabata and Kornberg 1994). Hh signaling does not occur in P compartment cells because they do not express critical components of the Hh pathway, such as the major transcriptional effector Ci (Eaton and Kornberg 1990). In contrast, A compartment cells can receive and respond to Hh but are unable to produce Hh. In A compartment cells located close to the source of Hh ligand production at the A/P boundary, Hh signaling triggers pathway activity and, consequently, an increase in the transcription of target genes (Ingham et al. 1991; Basler and Struhl 1994; Capdevila et al. 1994; Tabata and Kornberg 1994; Chen and Struhl 1996). A model of Hh secretion and transport from the basal surface of the *Drosophila* wing imaginal discs epithelia involves movement of Hh along cytonemes (Gradilla et al. 2014; Chen et al. 2017; Gonzalez-Mendez et al. 2017), which are dynamic thin cellular protrusions specialized for the intercellular exchange of signaling proteins (Ramirez-Weber and Kornberg 1999; Roy et al. 2011; Gradilla and Guerrero 2013; Kornberg 2014). Intriguingly, when Ihog is co-expressed with the cytoskeletal and membrane markers of these structures, these thin cellular protrusions are much easier to be detected microscopically (Callejo et al. 2011; Bilioni et al. 2013; Bischoff et al. 2013; Gonzalez-Mendez et al. 2017), which suggests that Ihog has roles in generating or stabilizing cytonemes. Moreover, overexpressed Ihog is used as a cytoneme marker to visualize these structures (Portela et al. 2019; Gonzalez-Mendez et al. 2020). Yet whether and how Ihog promotes cytoneme growth or stabilization and how cytonemes contribute to Hh transport and signal reception remain poorly understood.

Here, we report that cytoneme-localized Ihog proteins engage in *trans*-homophilic binding leading to cytoneme stabilization in a manner independent of the receptor role of Ihog in transducing the Hh signal. The Ihog-Ihog *trans*-homophilic binding site overlaps with the Ihog-Hh binding interface and requires the heparin binding site, suggesting direct competition between the dual roles of Ihog proteins. By combining experimental data and mathematical modeling, we determined Hh binding to Ihog dominates and can disrupt pre-established Ihog-Ihog *trans*-homophilic interactions, resulting in Hh-Ihog complexes. Our results indicated that the weaker Ihog-Ihog *trans* interactions promote and stabilize membrane contacts along the cytonemes and the disruption of some of these interactions by the stronger Hh-Ihog interaction could contribute to cytoneme-mediated transport of Hh or internalization of the ligand-receptor complex. Thus, we proposed that the apparently incompatible functions of Ihog in homophilic adhesion and ligand binding cooperate to assist Hh transport and reception along cytonemes.

## RESULTS

### Ihog stabilizes cytonemes in a manner independent of Hh receptor function

The Hh receptor component Ihog localizes to cytonemes in the *Drosophila* wing imaginal discs and abdominal histoblasts. Increasing Ihog abundance makes cytonemes in the histoblasts less dynamic and enables easier microscopic detection of these structures in the wing disc (Callejo et al. 2011; Bilioni et al. 2013; Bischoff et al. 2013; Gonzalez-Mendez et al. 2017). However, it is not clear how ectopic Ihog proteins influence the behavior and morphology of cytonemes. To explore the mechanism by which Ihog proteins affect cytonemes, we transiently expressed Ihog or the actin-binding domain of moesin fused to green fluorescent protein (GFP) (GMA-GFP) in the Hh-receiving cells in the A compartment using a *ptc-GAL4* driver in combination with *tub-GAL80*^ts^. Cytonemes projecting from the *ptc-GAL4* expressing cells were examined in the 3^rd^ instar larvae wing discs 24 hours after shifting to 29°C by staining with antibodies recognizing GFP or Ihog (Fig. 1A). Unlike the short, mostly uniform cytonemes visualized by staining for GMA-GFP, the cytonemes with ectopic expression of Ihog were longer with periodic dense structures (Fig. 1B, C). These dense structures are proposed to represent stable links between Hh-sending and Hh-receiving cytonemes (Gonzalez-Mendez et al. 2017).

**Figure 1.**
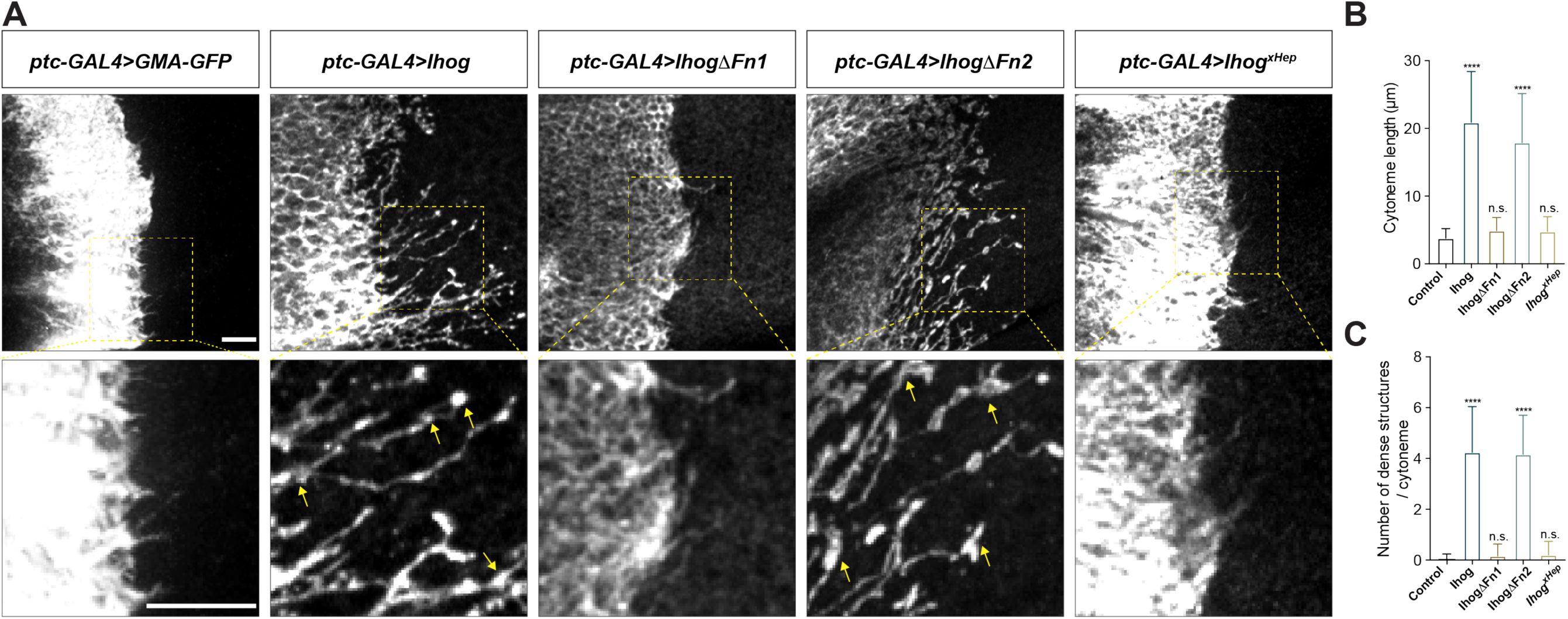
Ihog mediates cytoneme stabilization via the Fn1 domain. (A) Wing discs from 3^rd^ instar larvae carrying *ptc-GAL4, tub-GAL80^ts^* and the indicated *UAS-*transgenes were immunostained for GFP or Ihog to visualize cytonemes projecting from Hh-receiving cells. Yellow arrows indicate the dense structures along the cytonemes. Scale bar, 10 µm. (B, C) Quantification of the average cytoneme length (B) and the average dense structures number per cytoneme (C) in the wing disc. Each bar shows the mean ± SD (n > 30). One-way ANOVA followed by Dunnett’s multiple comparison test was used for statistical analysis. ns, not significant. ****P < 0.0001.

Several other Hh pathway components, including the Hh ligands, Ptc, and the *Drosophila* glypicans Division abnormally delayed (Dally) and Dally-like (Dlp), localize to cytonemes (Ayers et al. 2010; Chen et al. 2017; Gonzalez-Mendez et al. 2017). To determine if these effects of Ihog were unique to Ihog, we ectopically expressed each of these components individually using the *ptc-GAL4* driver in the wing imaginal disc cells. For these experiments, we included *UAS-Myr-RFP*, which encodes a myristoylated form of red fluorescent protein, to mark the cell membrane and enable visualization of the cytonemes. Of the tested Hh components, only expression of Ihog lead to formation of the dense structures or increased cytoneme length (Supplementary Fig. 1, Fig. 1A – C). We defined the increased cytoneme length and presence of dense structures as “cytoneme stabilization.”

Previous biochemical and structural studies showed that the Ihog Fn1 domain is involved in binding to the Hh ligand via a heparin-binding surface (McLellan et al. 2006; Yao et al. 2006), whereas the Fn2 of Ihog interacts with Ptc. Both Fn1 and Fn2 domains are required for formation of a high-affinity multimolecular complex of Ihog with Ptc and Hh during Hh signal reception (Zheng et al. 2010). We performed a structure-function analysis by expressing Ihog variants lacking either the first FNIII domain (Fn1) (IhogΔFn1) or the second Fn2 (IhogΔFn2) or with mutations in the heparin-binding surface (Ihog^xHep^) and quantified the cytoneme-stabilizing effects. These studies revealed that both the increased frequency of dense structures and length of the cytonemes required Fn1 and an intact heparin-binding surface (Fig. 1A – C).

*Ptc* not only encodes a component of the Hh receptor but is also a transcriptional target of Hh signaling. The highest expression of *Ptc* is in a stripe of A cells immediately adjacent to the A/P compartment boundary, whereas much lower *ptc* expression occurs in the A compartment away from the boundary and no expression occurs in the cells of the P compartment. We generated randomly distributed Ihog-overexpressing cells throughout the A and P compartments in the wing imaginal discs. We observed stabilized cytonemes emanating from with Ihog-expressing clones located not only at the Ptc^high^ A/P compartment boundary but also within the Ptc^low^ A compartment and Ptc^neg^ P compartment (Fig. 2A, upper row). Thus, the cytoneme-stabilizing effect of Ihog was independent of Ptc, consistent with the ability of Ihog lacking the Fn2 domain to perform this function. Moreover, the expression of Ptc-binding deficient IhogΔFn2 in the A/P boundary cells resulted in stable cytonemes projecting both posteriorly towards the Hh-secreting P cells and anteriorly away from the Hh source (Fig. 2B). These observations indicated that Ihog stabilizes cytonemes through a mechanism different from that used for the formation of Ihog-Ptc receptor complex, which exhibits high-affinity binding to Hh ligands.

**Figure 2.**
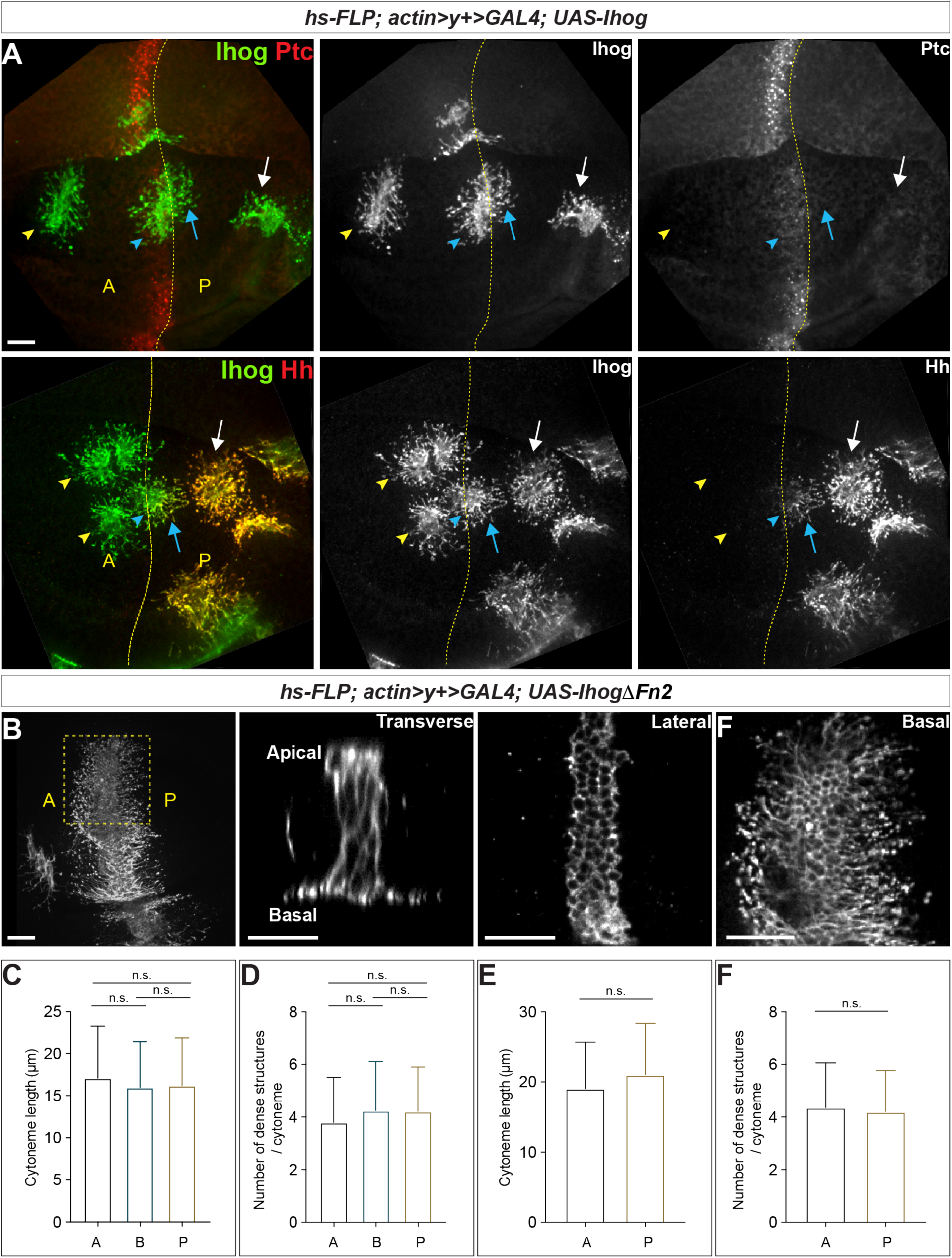
Ihog mediates cytoneme stabilization in a manner independent of the Hh receptor function. (A) Wing discs from 3^rd^ instar larvae carrying flip-out clones expressing *UAS-Ihog* were immunostained for Ihog (green) and Ptc or Hh (red) as indicated. Dashed yellow line indicate the A/P compartment boundary; white arrows indicate clones located within the P compartment; blue arrows indicate cytonemes of clones located at the A/P boundary that project toward the Hh producing cells; blue arrowheads indicate cytonemes of clones located at the A/P boundary that project away from the Hh producing cells; yellow arrowheads indicate cytonemes from clones located within the A compartment. Scale bar, 20 µm. (B) Flip-out clones expressing *UAS-IhogΔFn2* viewed from the basal side at low magnification and in transverse, lateral, and basal sections of the same clone, showing localization of IhogΔFn2 proteins at the apical-lateral/lateral cell-cell contacts and basal cytonemes. Scale bar, 20 µm. (C, D) Quantification of the average cytoneme length and the average dense structure number per cytoneme for A (clones located in the Ptc^low^ A compartment), B (Ptc^high^ A/P compartment boundary), and P (Ptc^neg^ P compartment). (E, F) Quantification of the average cytoneme length and the average dense structure number per cytoneme for A (cytonemes projecting anteriorly away from the Hh source) and P (cytonemes projecting posteriorly towards the source of Hh). Each bar shows the mean ± SD (n = 30). One-way ANOVA followed by Tukey’s multiple comparison test (C, D) or the two-tailed unpaired t-test (E, F) was used for statistical analysis. ns, not significant.

In the wing imaginal discs, Hh is secreted from the P compartment and there is little to no Hh that diffuses several cell diameters into the A compartment (Zecca et al. 1995; Mullor et al. 1997; Strigini and Cohen 1997). Ihog interacts with Hh both in the context of the Ihog-Ptc complex and independently from Ptc (McLellan et al. 2006; Yan et al. 2010; Zheng et al. 2010). We reasoned that, if Ihog Fn1-mediated binding to Hh contributes to the cytoneme-stabilizing effect, cytonemes projecting from the Ihog-expressing wing disc cells in the A or P compartment should display different properties. Consistent with Ihog interacting with Hh, Hh staining colocalized with Ihog-expressing cytonemes either projecting from clones located within the P compartment or from boundary clones projecting posteriorly toward the Hh source (Fig. 2A, lower row). No or very little Hh staining was detected with Ihog-expressing cytonemes from clones within the A compartment or with Ihog-expressing cytonemes from clones at the A/P boundary and extending into the A compartment. Despite the absence or limited amount of Hh ligands, Ihog expression stabilized all the cytonemes projecting within or toward the A compartment. These results indicated that Ihog Fn1-mediated binding to Hh ligands did not account for the stable cytonemes visualized by ectopic Ihog expression in the wing imaginal disc cells.

Quantification of cytoneme length and the number of dense structures per cytoneme for Ihog-expressing clones in the A compartment, P compartment, and at the boundary showed that cytoneme stabilization by Ihog was independent of position within the wing disc and thus the abundance of Ptc or Hh ligands (Fig. 2C, D). We also quantified cytoneme-stabilizing properties for IhogΔFn2 in flip-out clones located close to the A/P compartment boundary, which also showed no difference between cytonemes projecting posteriorly towards the Hh-secreting P cells and those projecting anteriorly away from the Hh source (Fig. 2E, F). Collectively, our results indicated that neither the presence of Ptc nor Hh is necessary for Ihog-mediated cytoneme stabilization. This function of Ihog was separate from its function in the Hh receptor.

### Ihog facilitates cytoneme stabilization through homophilic *trans* binding supported by glypicans

Previously, we showed that the Ihog Fn1 domain not only plays an essential role in Hh signal transduction but also mediates cell-cell interactions in a homophilic manner (Hsia et al. 2017; Wu et al. 2019). Because our data indicated that Ihog stabilized cytonemes through the Fn1 domain in a manner independent of Hh receptor function (Fig. 1, 2), we hypothesized that Ihog Fn1-mediated homophilic *trans* interactions were responsible for cytoneme stabilization. The region that we identified as mediating the *trans* Ihog-Ihog interaction overlaps with the region that mediates the interaction with Hh on the Ihog Fn1 domain and includes a region where the negatively charged glycan heparin binds (McLellan et al. 2006; Wu et al. 2019). *In vitro*, heparin is required for not only Ihog-Hh binding but also Ihog-Ihog homophilic *trans* interactions (McLellan et al. 2006; Wu et al. 2019). Thus, a model for Ihog-Ihog homophilic *trans* interactions involves heparin-dependent bridging of positively charged surfaces on the two opposing Fn1 domains, in a manner similar to heparin-bridged Ihog-Hh interactions (McLellan et al. 2006; Wu et al. 2019).

Heparin used in previous *in vitro* assays is an intracellular glycosaminoglycan (GAG) that is not present on the cell surface or along the cytonemes. Thus, heparin is unlikely to mediate Ihog-Ihog *trans* interactions *in vivo.* Heparan sulfate, which is an extracellular GAG structurally related to heparin and ubiquitously located on the cell surface or in the surrounding extracellular matrix, was subsequently found to supply the function of heparin and mediate Ihog-Hh interaction *in vitro* (Zhang et al. 2007). Heparan sulfate is also covalently attached to proteins forming heparan sulfate proteoglycans (HSPGs), thus, heparan sulfate or HSPGs may serve as the physiological correlate of heparin to mediate Ihog-Ihog homophilic *trans* binding and Ihog-Hh binding. Dally and Dlp are two *Drosophila* glycosylphosphatidylinositol (GPI)- anchored HSPGs, which can be membrane-tethered or released from cells upon cleavage (Bernfield et al. 1999). Dally and Dlp are known to be involved in modulating the transport and reception of the Hh signal (Lum et al. 2003; Lin 2004; Tabata and Takei 2004; Eugster et al. 2007; Ayers et al. 2010; Yan et al. 2010; Bilioni et al. 2013). Ihog-expressing cytonemes rarely extend across large clonal populations of *dally* and *dlp* double mutant cells (Gonzalez-Mendez et al. 2017), indicating that Dally and Dlp could be the major source of heparan sulfate that enables Ihog-Ihog homophilic *trans* interactions *in vivo*. Consistent with this hypothesis, we detected a striking accumulation of endogenous Dlp along Ihog-expressing cytonemes not only in the P compartment or along the A/P boundary where both Ihog-Hh and Ihog-Ihog interactions exist (Fig. 3A) but also within the A compartment that lacks Hh and where only Ihog-Ihog homophilic interactions could occur (Fig. 3B). Additionally, Dlp accumulated along the apical and lateral cell-cell contacts (Fig. 3B; yellow outlined regions), where homophilic Ihog *trans* binding contributes to cell segregation in the wing imaginal disc epithelium (Hsia et al. 2017; Wu et al. 2019). We also observed that ectopic expression of Ihog caused the accumulation of endogenous Dally along the apical and lateral cell-cell contacts and basal cytonemes (Supplementary Fig. 2).

**Figure 3.**
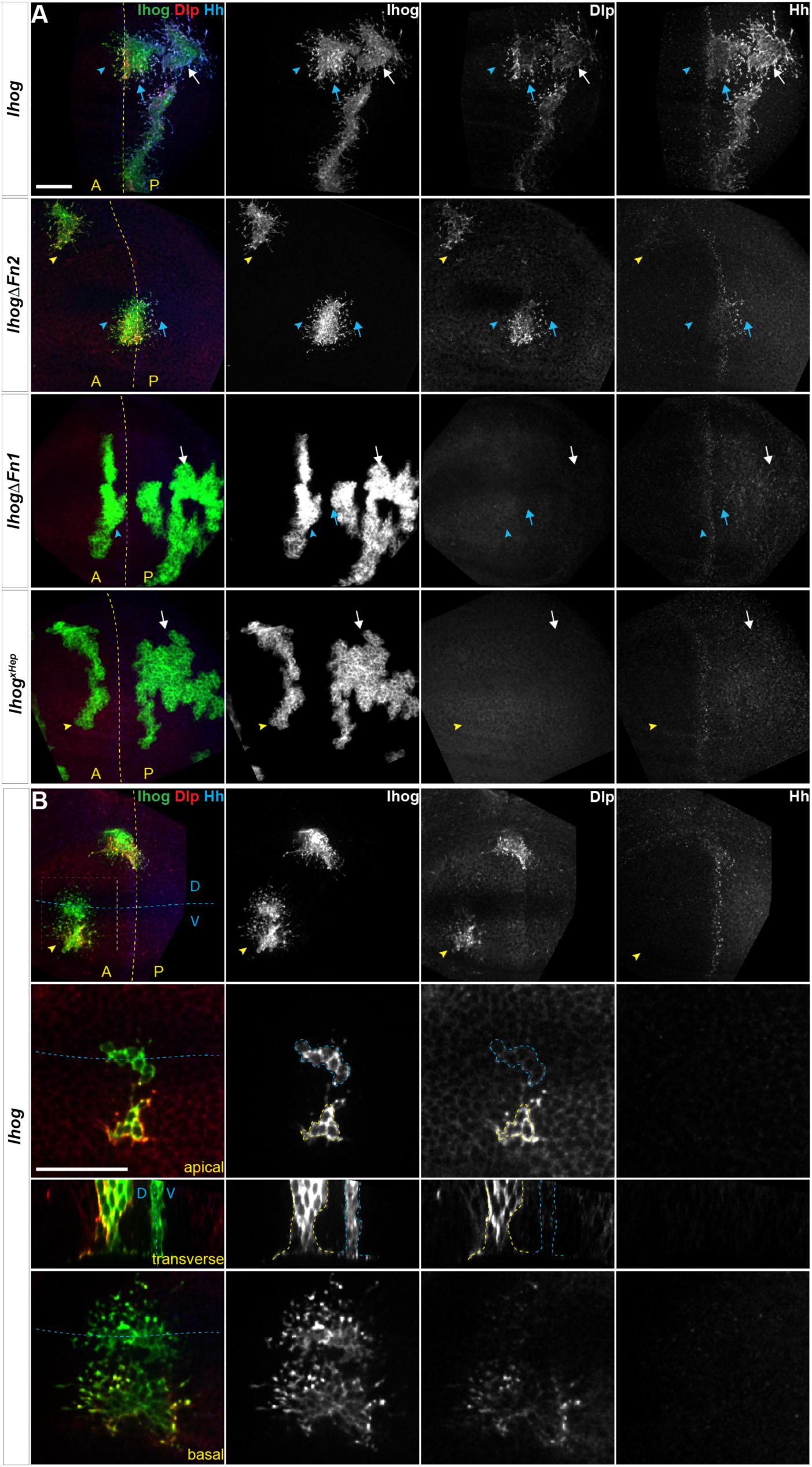
Ectopic Ihog induces accumulation of glypicans at lateral cell-cell contacts and along basal cytonemes. Wing discs from 3^rd^ instar larvae carrying flip-out clones expressing the indicated *UAS-* transgene were immunostained for Ihog (green), Dlp (red), and Hh (blue). Dashed yellow lines indicate the A/P compartment boundary; dashed blue lines indicates the dorsal/ventral (D/V) compartment boundary. (A) Ihog or Ihog mutants were expressed in wing discs. White arrows indicate clones located within the P compartment; blue arrows indicate cytonemes of clones located at the A/P boundary that project toward the Hh producing cells; blue arrowheads indicate cytonemes of clones located at the A/P boundary that project away from the Hh producing cells; yellow arrowheads indicate cytonemes from clones located within the A compartment. (B) Flip-out clones expressing *UAS-Ihog* viewed from the basal side at low magnification and in lateral, transverse, and basal sections of the zoomed area. The upper row shows the imaged clones (outlined with dashed white lines) with their position relative to the A/P and D/V boundaries. Blue outline indicates the flip-out clone flanking the D/V boundary; the yellow outline indicates the clone several cell diameters away from the D/V boundary. Scale bar, 20 µm.

The Ihog-induced accumulation of Dally and Dlp along the basal cytonemes is consistent with a crucial contribution of either Dally or Dlp in Ihog-mediated cytoneme stabilization (Gonzalez-Mendez et al. 2017). However, ectopic expression of neither Dally nor Dlp resulted in cytoneme stabilization in the wing discs (Supplementary Fig. 1). Thus, the roles of Ihog and the glypicans in stabilizing the cytonemes are different. Given the known function of heparin as a bridging molecule in Ihog-Ihog or Ihog-Hh interactions, our results suggested that the heparan sulfate chains of Dally or Dlp provide this function in Ihog-Ihog *trans* interactions. Consistent with this model, the Ihog-induced Dally or Dlp accumulation reflected the different distributions of the two glypicans. Dlp is distributed in most cells, except in a zone ∼7 – 10 cell diameters in width and centered at the dorsal ventral (D/V) boundary. Dally is also broadly distributed; however, Dally abundance is highest along the D/V boundary (Fujise et al. 2001; Fujise et al. 2003; Han et al. 2005). In agreement with these different distributions, both Dally and Dlp were enriched along the cytonemes and at the apical and lateral cell contacts of Ihog-expressing cells away from the D/V boundary, whereas the Ihog-expressing cells flanking the D/V boundary were positive for Dally with little or no detectable Dlp (Fig. 3B, blue outline; Supplementary Fig. 2). Similar to the Ihog-mediated homophilic binding, Ihog-induced Dlp accumulation occurred in the absence of the Ihog Fn2 domain (Fig. 3A). In contrast, neither ectopic expression of IhogΔFn1 nor Ihog^xHep^, both of which lack homophilic binding capability, resulted in the accumulation of Dlp (Fig. 3A). Taken together, these results indicated that Ihog Fn1-mediated homophilic *trans* interactions, assisted by the heparan sulfate chains of Dally or Dlp in the wing imaginal discs, contribute to cytoneme stabilization.

### Homophilic Ihog *trans* interactions promote direct cytoneme-cytoneme contact formation

We previously found that ectopic expression of Ihog in the non-adherent *Drosophila* S2 cells induces cell aggregation via homophilic *trans*-interactions (Wu et al. 2019). Pairs of Ihog-positive S2 cells in close contact (aggregated) showed a peak of Ihog positivity along the site of cell-cell contact (Supplementary Fig. 3A-C, Supplementary movies 1, 2). In contrast, Ihog was enriched in filopodia-like structures on dispersed S2 cells when evaluated for live cells or fixed cells (Supplementary Fig. 3D-G, Supplementary movies 3, 4). Because *Drosophila* S2 cell filopodia recapitulate structural and functional characteristics of cytonemes in the imaginal disc (Bodeen et al. 2017), here, we take advantage of these dispersed Ihog-expressing S2 cells to evaluate the possibility of filopodia-localized Ihog proteins in participating homophilic *trans*-interactions. We examined the behavior of these Ihog-positive filopodia between an Ihog-positive cell and an Ihog-negative cell and between pairs of Ihog-positive cells. These structures were found at regions where two Ihog-positive cells were in close apposition with the filopodia projecting toward the adjacent Ihog-positive cell (Supplementary Fig. 4A). In contrast, an Ihog-positive cell extend fewer filopodia toward an Ihog-negative cell (Supplementary Fig. 4A, B). Two closely positioned Ihog-positive cells exhibited an increase in the number of filopodia oriented toward the nearby Ihog-positive cell (Supplementary Fig. 4C). Moreover, using live-cell time-lapse imaging, we observed that the nearby cells that were both positive for Ihog became closer as the Ihog-positive filopodia became more interdigitated (Supplementary Fig. 5). Collectively, the observations in S2 cells suggested that filopodia-localized Ihog proteins engaged in homophilic *trans* binding evidenced by the contact initiation among non-adjacent Ihog-expressing S2 cells.

To explore if such events occurred *in vivo*, we generated Ihog-expressing clones in the wing imaginal discs and found that cytonemes projecting from closely positioned clones appeared to come into contact (Figs. 4A; arrows). Unlike cultured S2 cells, the wing imaginal disc epithelial cells tightly adhere to their immediate neighbors and maintained stable cell-neighbor relationships (Garcia-Bellido et al. 1973; Gibson et al. 2006). Cytoneme-cytoneme interaction is unlikely to lead to move the cell body, thus a reduction in the distance between non-adjacent Ihog-expressing clones could not be used as the functional readout of Ihog-Ihog *trans*-interaction along the cytonemes. Therefore, we examined whether direct membrane contacts were established along Ihog-localized cytonemes.

Membrane contacts are typically separated by less than 100 nm of extracellular space, which is below the resolution of conventional light microscopy. To examine whether membrane contacts were established along Ihog-stabilized cytonemes, we combined the CoinFLP-LexGAD/GAL4 system and the GFP Reconstitution Across Synaptic Partners (GRASP) system (Feinberg et al. 2008; Gordon and Scott 2009; Bosch et al. 2015). CoinFLP-LexGAD/GAL4 allows generation of tissues composed of clones that express either GAL4 or LexGAD, thus enabling the study of interactions between different groups of genetically manipulated cells (Bosch et al. 2015). In the GRASP system, two complementary parts of a ‘split GFP’ (spGFP_1-10_ and spGFP_11_) are fused to the extracellular domains of mouse CD4, one under UAS control and the other under LexAop control. Whereas individually the membrane-tethered spGFP fragments are not fluorescent, reconstitution of GFP generates fluorescence at the boundary of immediately adjacent clones that express the complementary spGFP fragments (Bosch et al. 2015). We expressed RFP-tagged Ihog together with CD4-spGFP_1-10_ and HA-tagged IhogΔFn2 with CD4-spGFP_11_. With this system, we monitored cells for the presence of either Ihog or IhogΔFn2 using an antibody recognizing Ihog and cells positive for only IhogΔFn2 using an antibody against the HA tag.

As expected from the CoinFLP system, clones expressing Ihog-RFP plus CD4-spGFP_1-10_ and those expressing IhogΔFn2-HA plus CD4-spGFP_11_ were randomly distributed in the wing imaginal discs. When these two types of clones were located immediately adjacent to each other, GRASP fluorescence was detected both at the apical lateral contacts and along the basal cytonemes of the adjacent cells expressing Ihog-RFP plus UAS-CD4-spGFP_1-10_ or IhogΔFn2-HA plus LexAop-CD4-spGFP_11_ (Fig. 4B, C, D; blue outline and blue arrow). The basal cytonemes emanating from wing imaginal disc cells can reach as far as several cell diameters, thus, if cytoneme-localized Ihog proteins participate in homophilic *trans*-binding, direct cytoneme-cytoneme contacts from non-adjacent cells could be preferentially established among these cytonemes expressing ectopic Ihog proteins. Indeed, GRASP fluorescence also appeared along the length of the cytonemes projecting from the non-adjacent Ihog-RFP plus UAS-CD4-spGFP_1-10_ and IhogΔFn2-HA plus LexAop-CD4-spGFP_11_ expressing cells that do not share common boundaries as indicated by lacking of GRASP florescence throughout the apical and lateral clonal borders (Fig. 4B, C, D; yellow outline and yellow arrow). Therefore, the GRASP marked cytoneme-cytoneme contacts from non-adjacent clones suggested that membrane contacts were initiated and established along Ihog-expressing cytonemes, supporting the idea that Ihog-Ihog *trans*-binding can occur along opposing cytoneme membranes *in vivo*.

**Figure 4.**
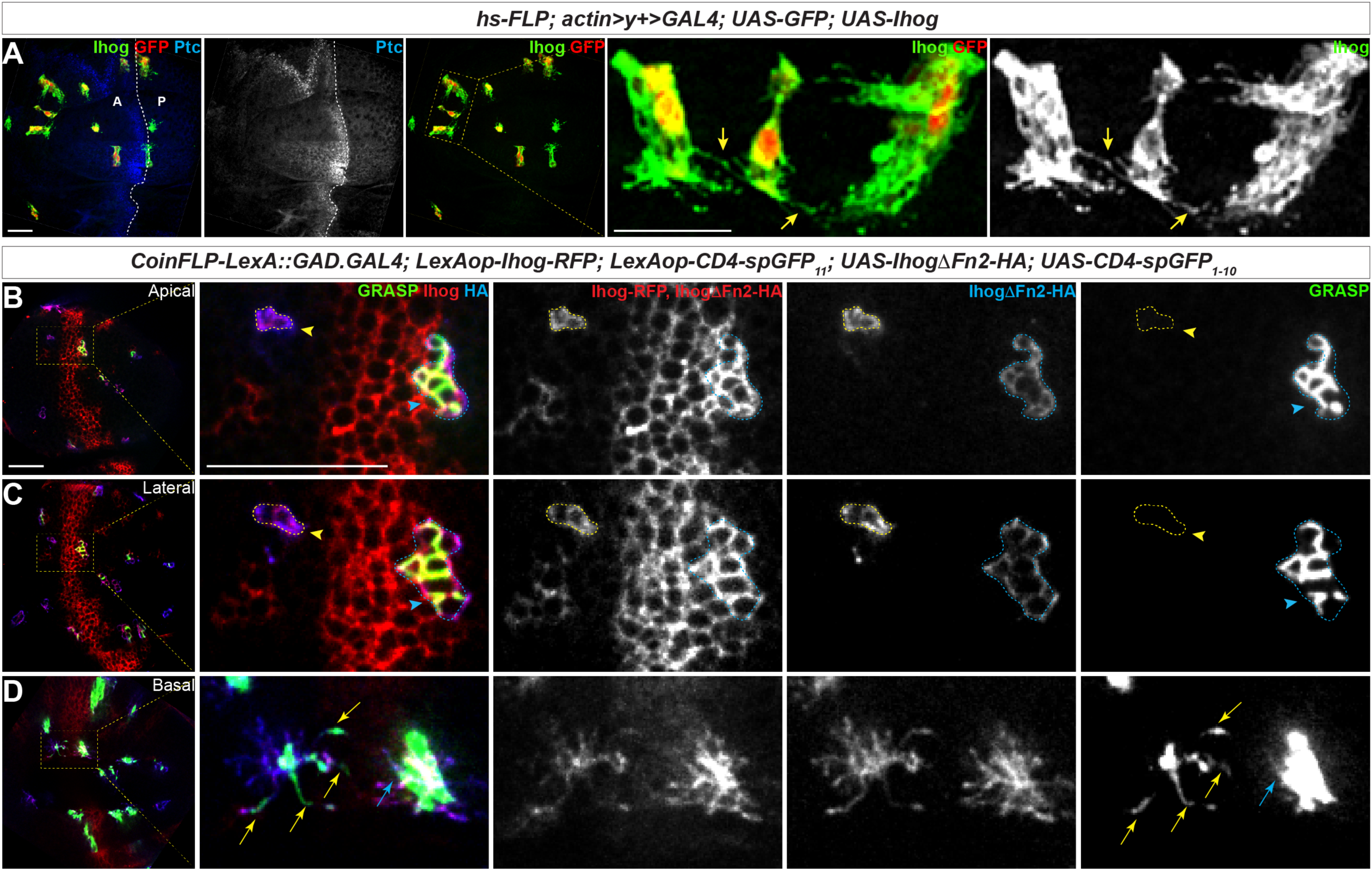
Homophilic Ihog *trans* binding enables direct cytoneme-cytoneme contact formation. (A) Wing imaginal discs from 3^rd^ instar larvae carrying flip-out clones expressing *UAS-Ihog* were immunostained with antibodies against Ihog (green), Ptc (blue), and GFP (red) as indicated. Yellow arrows indicate Ihog-enriched cytonemes projecting from closely positioned clones. Scale bar, 20 µm. (B-D) Apical, lateral, and basal sections of a wing imaginal disc from 3^rd^ instar larvae carrying clones marked by the CoinFLP-LexGAD/GAL4 system and the GRASP system as indicated. The wing discs were immunostained with antibodies against Ihog (red, both Ihog-RFP and IhogΔFn2-HA expressing cells) and HA (blue, IhogΔFn2-HA expressing cells) as indicated. GRASP signal is green. Blue outlines indicate clones expressing CD4-spGFP_1-10_ and IhogΔFn2-HA that are immediately adjacent to clones expressing CD4-spGFP_11_ and Ihog-RFP; yellow outlines indicate CD4-spGFP1-10 and UAS-IhogΔFn2-HA clones that are distant from those expressing CD4-spGFP_11_ and Ihog-RFP. Blue arrowheads indicate GFP fluorescence along the apical and lateral sides of the outlined clones. Yellow arrowheads indicate absence of GFP fluorescence along the apical and lateral sides of the outlined clones. (D) Blue and yellow arrows indicate GFP fluorescence along the length of the cytonemes projecting from the clones indicated by blue and yellow outlines (B, C), respectively. Scale bar, 20 µm.

### Modeling predicts that homophilic Ihog *trans* interactions increase cytoneme length and bundling

We developed a stochastic model to investigate the influence of the homophilic *trans*-interaction strength on the dynamics of cytonemes. In the model, cytonemes were represented as filamentous structures with variable numbers of discrete segments. We considered the elongation, shrinkage, translocation, and interaction events of cytonemes around the cell surface: An elongation or shrinkage event was represented by the addition or removal of one segment to or from an existing cytoneme; a translocation event was represented as the movement of a cytoneme along the cell surface; and interaction events of two cytonemes in contact were represented by pairwise interactions between segments.

We set the elongation probability of the *i^th^* cytoneme to exponentially decay with its length:

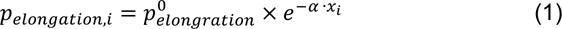

where 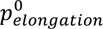 is the base elongation probability at the cell surface, *α* is the decay coefficient, and *x_i_* is the number of segments in the *i^th^* cytoneme (that is the length that the cytoneme extended from the cell surface). This decay relationship represents the increasing difficulty in transporting materials to the tip of the cytoneme as elongation occurs and increasing difficulty in the occurrence of elongation as the membrane tension increases, thereby resisting elongation.

The shrinkage rate at the tip of the *i^th^* cytoneme is modeled as

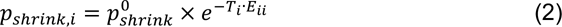

where 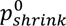 is the intrinsic shrinkage rate without any homophilic *trans* interaction, *E_ii_* > 0 is the strength of the homophilic *trans* interaction between a pair of segments, and *T_i_* is the number of neighboring cytonemes with which this *i^th^* tip segment interacts. Thus, the homophilic *trans* interactions at the tip segment represent additional energy barriers to a shrinkage event.

To enhance simulation efficiency, we employed the quasi-equilibrium approximation (Goutsias 2005) to simulate the pairwise interactions between segments on neighboring cytonemes. First, we computed the probability of establishing a homophilic *trans* interaction between a pair of neighboring cytoneme segments:

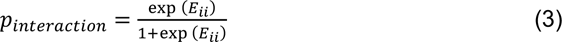

Based on *p_interaction_*, we randomly assigned a binary state variable, 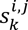 (0 for not interacting, 1 for interacting), to the *k^th^* pair of neighboring segments in the *i^th^* and *j^th^* cytonemes for each simulation step. Thus, the total homophilic interactions of the *i^th^* cytoneme is calculated as

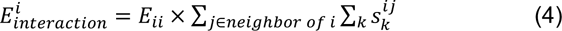

The cytonemes can also translocate along the cell periphery. A translocation event involves breaking the existing homophilic *trans* interactions and establishing new homophilic *trans* interactions. We computed the differences in the *trans* interactions before and after a possible translocation event for the *i^th^* cytoneme as 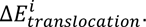 Using the computed probabilities and energy differences, we performed stochastic simulations and collected the cytoneme configurations after the simulations reached steady state (see Methods for details).

We simulated the system with no (*E_ii_* = 0) homophilic *trans* interactions (Fig. 5A, left) and with homophilic *trans* interactions of moderate strength (*E_ii_* = 15) (Fig. 5A, right). Without any homophilic *trans* interactions, the simulations resulted in much fewer numbers of segments (shorter cytoneme length). For *E_ii_* = 15, the simulations predicted more variability in the length of cytonemes than was predicted at *E_ii_* = 0. By capturing 1001 snapshots from the random simulations for *E_ii_* = 0 and 15, we found that the simulations produced cytoneme lengths that were significantly longer at *E_ii_* = 15 (Fig. 5B). Additionally, the number of established pairwise interactions between cytonemes greatly increased at *E_ii_* = 15 (Fig. 5C). Thus, the simulations indicated that cytoneme length correlated with the number of cytoneme-cytoneme interaction events (Fig. 5D, Pearson r = 0.7939). By varying the strength of homophilic *trans* interactions, we also found that average cytoneme length increased with stronger cytoneme-cytoneme interactions (Fig. 5E).

**Figure 5.**
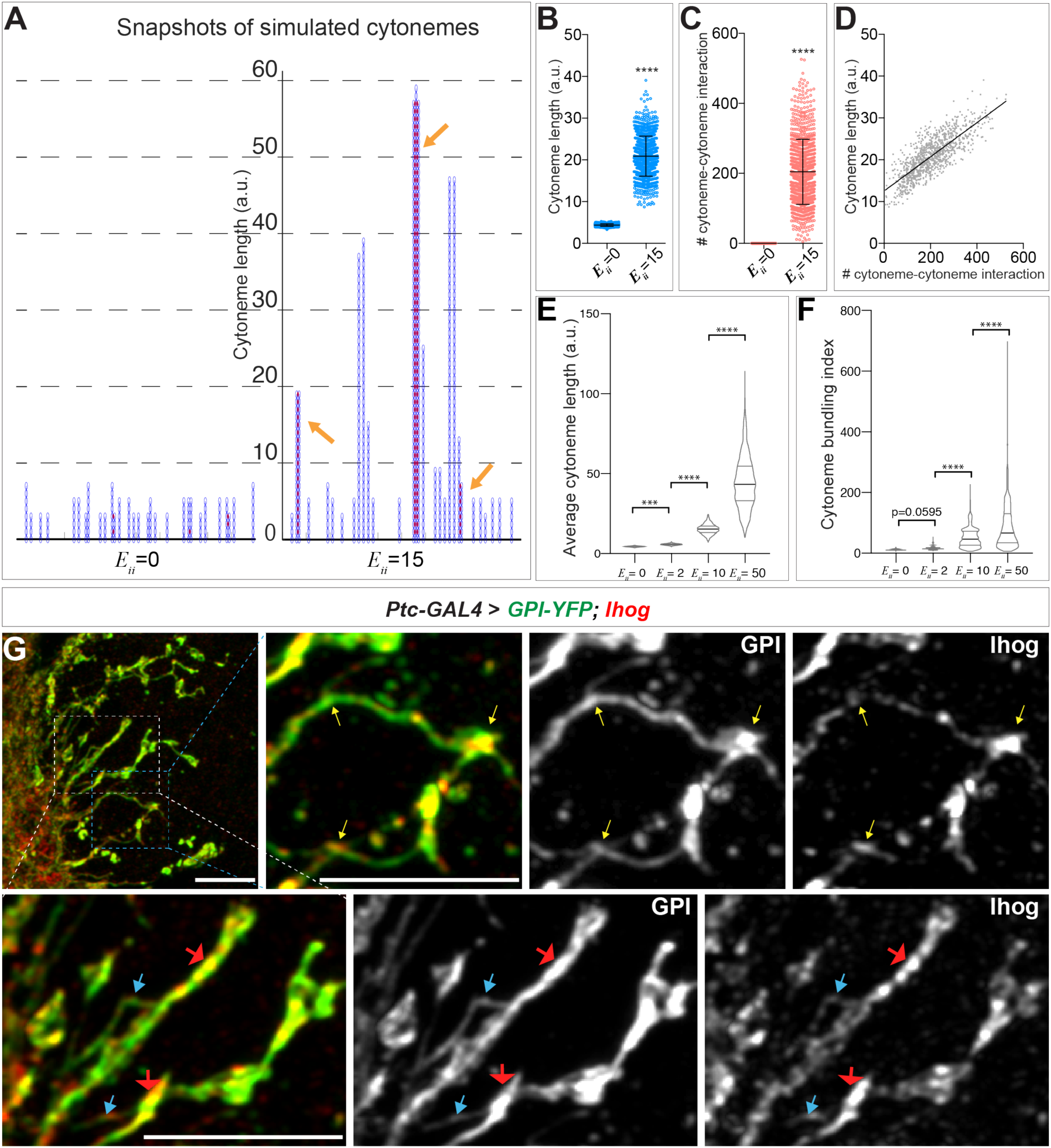
Computational modeling predicts that homophilic *trans* interactions stabilize cytonemes. (A) Snapshots of simulated cytoneme configurations with no (left, *E_ii_* = 0) homophilic *trans* interactions and with moderately strong (right, *E_ii_* = 15) homophilic *trans* interactions. The solid black horizontal lines at the bottom represent the cell surface. The blue vertical filaments are cytonemes, within which the elliptical elements are the individual segments. The red dots are the established pairwise interactions between neighboring segments. The orange arrows indicate the bundled neighboring cytonemes with extensive pairwise contacts. (B, C) The average cytoneme length and the number of cytoneme-cytoneme interactions at *E_ii_* = 0 and 15. Each dot is obtained from 1 randomly picked snapshot from the simulation. Each bar shows the mean ± SD, n = 1001. (D) Correlation between the cytoneme length and the number of pairwise interactions at *E_ii_* = 15. Each dot represents the length and the number of interactions for individual cytonemes. The grey line shows the best fit linear regression (Pearson r = 0.7939). (E) Effect of homophilic *trans*-interaction strength on the average cytoneme length. The average length of the simulated cytonemes is plotted against *E_ii_* ranging from 0 to 50. Each bar shows the mean ± SD, n = 1001. (F) Effect of homophilic *trans*-interaction strength on the formation of cytoneme bundles. The cytoneme bundling index for all cytoneme bundles identified from n=1001 random snapshots, each containing 30 cytonemes, were plotted against *E_ii_* ranging from 0 to 50. Each bar shows the mean ± SD, n = 1001. The two-tailed unpaired t-test (B, C) or one-way ANOVA followed by Sidak’s multiple comparison test (E, F) was used for statistical analysis. ***P < 0.001, ****P < 0.0001. (G) Wing discs from 3^rd^ instar larvae carrying *ptc-GAL4, tub-GAL80^ts^* and *UAS-GPI-YFP; UAS-Ihog-RFP* were immunostained for YFP (GPI, green) and Ihog (Ihog, red), followed by imaging with Airyscan. Yellow arrows indicate cytoneme-cytoneme contacts; blue arrows indicate likely single cytonemes; red arrows indicate bundles containing multiple cytonemes. Scale bar, 5 µm.

In the snapshots of the simulations, we observed extensive pairwise interactions among adjacent cytonemes only when we set *E_ii_* > 0 (Figs. 5A, arrows). We defined this phenomenon as “cytoneme bundling” and quantified this phenomenon with a cytoneme bundling index (see Methods). The cytoneme bundling index increased as the strength of the homophilic *trans* interactions among cytonemes increased (Fig. 5F). Furthermore, with increasing cytoneme-cytoneme homophilic *trans*-interaction strength *E_ii_*, we observed a decreased proportion of singular cytonemes and an increased proportion of cytonemes within the cytoneme bundles (Supplementary Fig. 6). These results predicted that cytonemes of Ihog-overexpressing cells would form extensive contacts and appear as bundles. By regular confocal microscopy, we observed an increase in dense contact sites, but we did not detect clear evidence of cytoneme bundles in the Ihog-overexpressing wing disc. This is likely because the diameter of cytonemes [100–200 nm (Mattila and Lappalainen 2008; Kornberg 2014)] is much less than the resolution limit (∼250 nm laterally) of confocal microscopy. We used Airyscan technology, which has a lateral resolution of 120 nm (Huff 2015)., to test the prediction of Ihog-induced bundling of cytonemes in the wing disc. We imaged Ihog-expressing cytonemes in wing discs cells co-labeled with membrane marker glycosyl-phosphatidyl-inositol–YFP (Greco et al. 2001) (GPI– YFP, Fig. 5G). Consistent with the computational modeling prediction, we detected sites of cytoneme-cytoneme contact (Fig. 5G, yellow arrows) and thin cytonemes (Fig. 5G, blue arrows) that appeared to form thick bundles (Fig. 5G, red arrows). In contrast, we rarely observed cytoneme bundling upon expression of only the membrane marker GPI-YFP or expression of the homophilic binding-deficient Ihog variants IhogΔFn1 and Ihog^xHep^ (Supplementary Fig. 7). Also consistent with the model predictions, we found that knockdown of *ihog* in the absence of its close paralog-encoding gene *boi* resulted in cytonemes with significantly reduced length compared with the length of cytonemes in *boi* mutant animals retaining normal expression of Ihog (Supplementary Fig. 8). Thus, the *in vivo* observations supported the predictions from the computational modeling that the augmented cytoneme-cytoneme interactions mediated by ectopic Ihog lead to elongated and bundled cytonemes.

### Heterophilic Ihog-Hh binding dominates over homophilic Ihog *trans* interaction

A single Ihog protein can participate in either an Ihog-Ihog *trans* interaction or an Ihog-Hh interaction; therefore, a single Ihog protein can mediate either its cytoneme-stabilizing function or its ligand-binding function, but not both simultaneously. The dissociation constant for the nonlipid-modified recombinant HhN and the extracellular portion of Ihog containing the Fn1 and Fn2 domains (IhogFn1-2), measured in solution, is ∼ 2 μM (McLellan et al. 2006; Zhang et al. 2007). Additionally, soluble HhN at 30 μM competes for Ihog homophilic interactions in a pull-down assay in which HhN and Ihog are mixed simultaneously (Wu et al. 2019). To explore the interaction hierarchy of Ihog-Ihog *trans* interactions and Ihog-Hh interactions, we used the S2 cell system. Ihog-Ihog *trans* interactions result in aggregation of the normally non-adhesive S2 cells (Hsia et al. 2017; Wu et al. 2019). We performed Ihog-mediated cell aggregation assays with two populations of cells, one expressing Ihog and GFP and the other expressing Ihog and mCherry, in the presence of exogenously applied recombinant HhN. Unexpectedly, even at 30 µM, a concentration 10 times higher than the reported dissociation constant, soluble HhN had little effect on Ihog-mediated cell aggregation (Supplementary Fig. 9). One explanation is that soluble HhN does not reach a sufficiently high concentration at the cell surface to compete for the extensive homophilic *trans* interactions that can be mediated by membrane-tethered Ihog proteins. Thus, we developed a system to test the ability of plasma membrane-associated Hh to interfere with Ihog-Ihog *trans* interactions. Expression of cDNA encoding full-length Hh in S2 cells generates dually lipid-modified Hh ligands, whereas expressing cDNA encoding the amino-terminal signaling fragment results in HhN lacking a cholesterol modification (Burke et al., 1999; Porter et al., 1995). Although both forms of Hh ligands are competent to bind to the receptors and induce Hh signaling in ligand-receiving cells, HhN does not require Dispatched (Disp) for release from the producing cell. Thus, in S2 cells without also ectopically expressing Disp, only dually lipid-modified Hh ligands resulted from cDNA encoding the full-length Hh and this lipid-modified Hh was enriched at the surface of the transfected cells (Supplementary Fig. 10). Using these cells, we assessed the relative strengths of Ihog-mediated ligand binding and homophilic *trans* interactions.

A heterogeneous aggregation of cells exhibits distinct morphological patterns when the relative strengths of the heterotypic and homotypic cell-cell adhesions differ. For example, a checkerboard-like pattern can occur when heterotypic cell-cell adhesions dominate (Honda et al. 1986). Therefore, we hypothesized that the morphological patterns produced by the aggregated Hh- and Ihog-expressing cells reflect the relative affinity of Ihog-Hh (heterotypic) and Ihog-Ihog (homotypic) interactions. To test this hypothesis, we prepared S2 cells expressing Hh or HhN and cells expressing Ihog, along with either GFP or monomeric Cherry (mCherry), mixed the cells, and assessed the pattern of the aggregated clusters (Fig. 6). We found that Hh-expressing cells remained dispersed when cultured by themselves (Fig. 6A, mCherry+Hh, GFP+Hh) and aggregated when mixed with Ihog-expressing cells (Fig. 6A, mCherry+Ihog, GFP+Hh). In contrast, HhN-expressing cells remained dispersed when cultured by themselves (Fig. 6A, mCherry+HhN, GFP+HhN).

**Figure 6.**
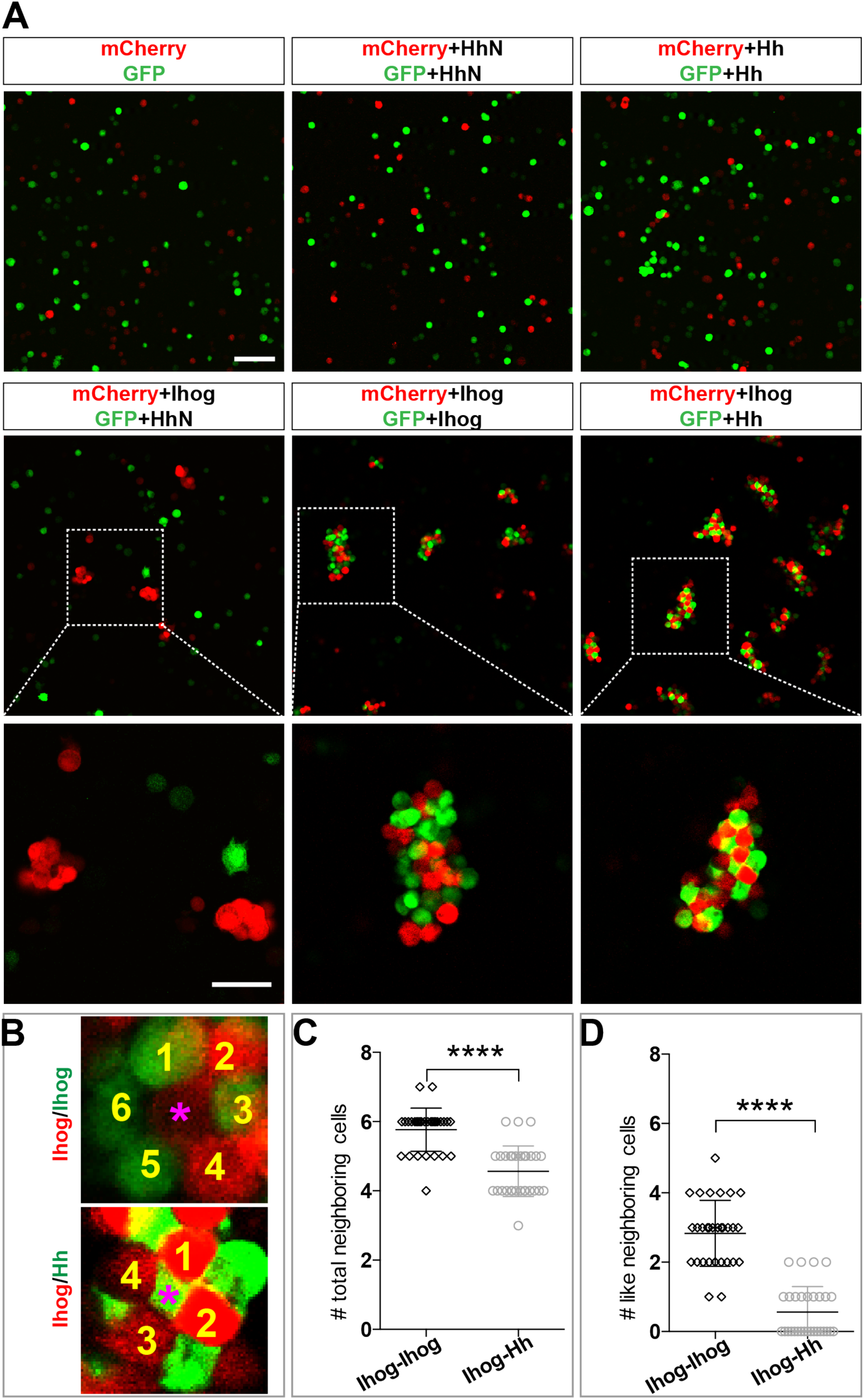
Heterophilic binding of Ihog to Hh dominates over Ihog-mediated homophilic *trans* interactions. (A) S2 cells were transfected with plasmids expressing Ihog, Hh, or HhN along with expression plasmids for either GFP or mCherry as indicated. Cells were dissociated by trypsin treatment and then mixed for 12 hr to allow aggregation to occur. The top and middle rows show the mixing of cells expressing only the fluorescent proteins or together with Hh, HhN or Ihog. Scale bar, 100 µm. The bottom row shows the indicated zoomed area from the middle row images. Scale bar, 50 µm. (B) Representative examples used for quantification of cell patterns in aggregates. The center cells in cell aggregates are indicated by purple asterisks. The neighboring cells are counted and labeled with yellow numbers. (C, D) The average numbers of total neighbor cells and “like” (expressing the same proteins and thus the same color as the center cell) neighbor cells were quantified. Each bar shows the mean ± SD, n = 30 cells (from n>3 experiments). The unpaired two-tailed t-test was used for statistical analysis. ****P < 0.0001.

When HhN-expressing cells were cocultured with Ihog-expressing cells, only the Ihog-expressing cells clustered (Fig. 6A, mCherry+Ihog, GFP+HhN). We compared the patterns of the cells in the clusters containing only Ihog-expressing cells and those containing both Ihog-expressing cells and Hh-expressing cells. We observed a checkerboard-like pattern with evenly distributed red and green cells in aggregates formed by cells expressing Hh with GFP and cells expressing Ihog with mCherry (Fig. 6B). Most center cells within the cell aggregates had 4 or 5 neighbors (Fig. 6C). Furthermore, among those neighbors, cells expressing the same transfected proteins and thus of the same color (“like” cell) were rare (Fig. 6B, D). In contrast, aggregates of Ihog-expressing cells labeled with either GFP or mCherry exhibited a honeycomb pattern (Fig. 6B): Most center cells had 5 or 6 neighbors (Fig. 6C), and ∼50% were “like” cells (Fig. 6B, D). For each aggregation assay, we confirmed by immunoblotting that transfected cells from the same experiment used for the aggregation assays expressed comparable amounts of Ihog and Hh proteins (Supplementary Fig. 11). Therefore, the different cellular patterns formed by cells expressing Hh and cells expressing Ihog versus those formed by differentially labeled Ihog-expressing cells suggested that the heterophilic interaction between Ihog and Hh is stronger than the homophilic *trans* interaction between Ihog proteins on an opposing cell surface.

### Computational modeling estimates the difference in strength between the heterophilic Ihog-Hh and homophilic Ihog *trans* interaction

Directly determining the affinities of the homophilic and heterophilic interactions is difficult because the affinities depend on the conformation of the Ihog dimers (*trans*-interacting or *cis*-interacting) and the membrane association of Hh. Thus, we took a computational approach to estimate the relative affinities of these two Ihog interactions. Motivated by the observations that cells expressing Hh and Ihog produced a different pattern from the cells expressing Ihog, we estimated the difference in strength between the heterophilic Ihog-Hh and homophilic Ihog-Ihog *trans* interactions by modeling these interactions using a vertex-based *in silico* assay (Bi et al. 2015; Park et al. 2016). We explicitly included heterogeneous cell composition in our model in the following manner: The cells were approximated by polygons that can freely change their locations and shapes. Consequently, two interacting cells were represented by two polygons sharing a common edge. This interaction leads to an energy reduction, the magnitude of which depends on various properties including the strength of the cell-cell adhesive interactions. From the cell shapes and configurations of neighboring cells, mechanical energy (*e_i_*) was calculated for each cell according to (Farhadifar et al. 2007) as:

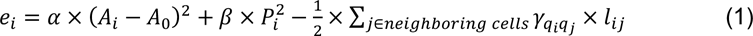

The first term is the areal elasticity, which is represented by *α* (the elastic coefficient), *A_i_* (the area of the *i*th cell), and *A*_0_ (the preferred cell area). The second term is the contractile energy, which is represented by *β* (the contractile coefficient) and *P*_+_ (the perimeter of the *i*th cell). The third term is the net adhesive energy between the *i*th cell and its neighbors, where *γ_qiqj_* is the line density of the adhesive energy between cell types *q_i_* and *q_k_*, and *l_ij_* is the length of the common edge between the two cells. We have *γ γ_II_*, *γ_IH_*, and *γ_HH_*, depending on the types of surface proteins expressed by the cells: both expressing Ihog, *II*, or one expressing Ihog and one expressing Hh, *IH*. Here, *γ*_HH_ = 0, because we did not observe cells both expressing Hh in contact with each other in the aggregation assays.

We used the Monte Carlo method (Metropolis 1953) to simultaneously simulate 100 cells within a 2-dimensional space. Gaps between cells were simulated as empty polygons that do not contribute to the mechanical energy of the system. With this system, the aggregation or segregation of cells is governed by ∑ *e_i_*. As a control, we simulated 100 Ihog-expressing cells with half labeled red and half labeled green, which produced a honeycomb morphological pattern (Fig. 7A), within which a given center cell had 5.8 ± 0.6 neighbors, and 2.4 ± 0.9 of them had the same color label as the center cell (Fig. 7C, I∼I bars).

**Figure 7.**
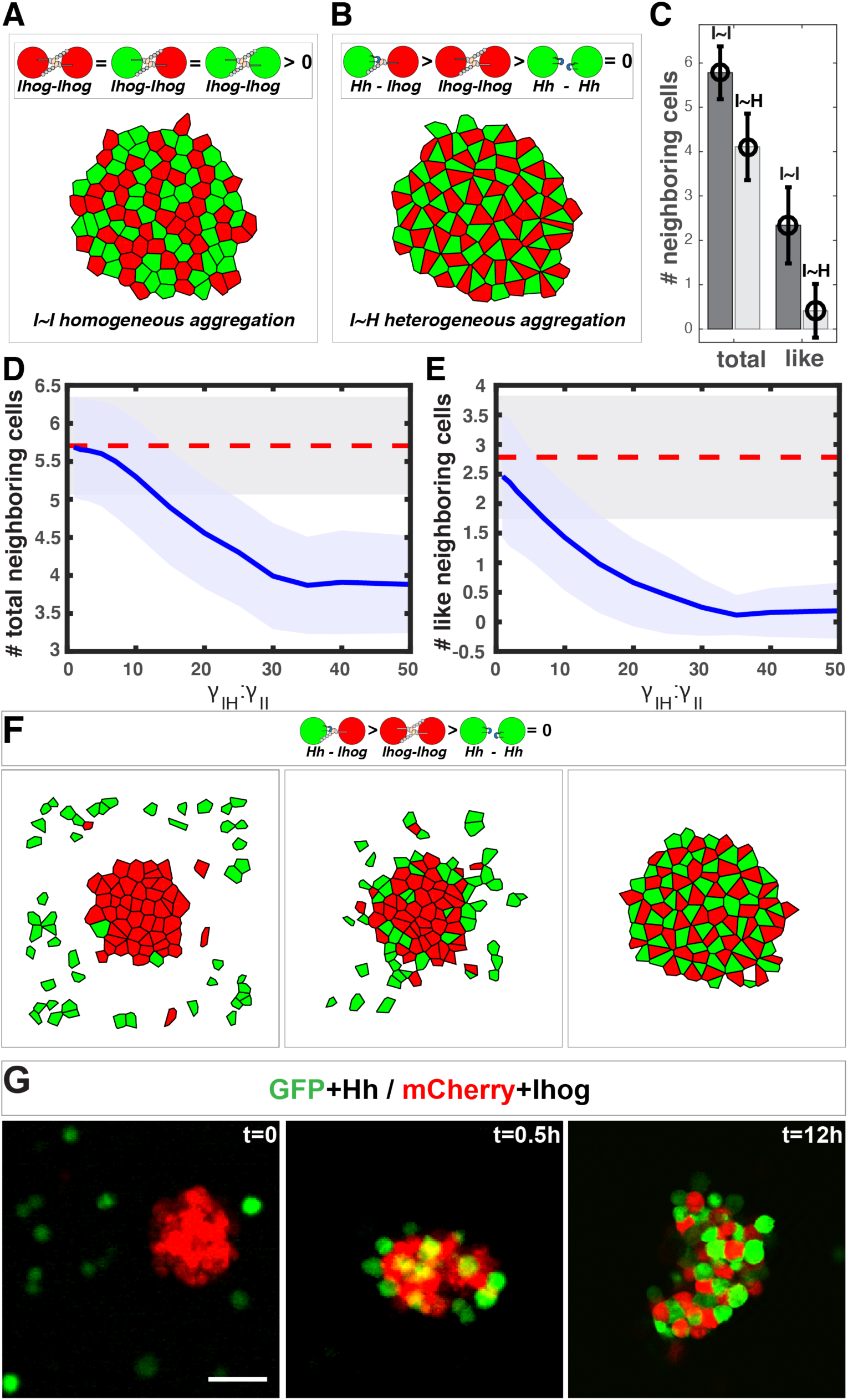
*In silico* simulation estimates the difference in strength between the heterophilic Ihog-Hh and homophilic Ihog-Ihog *trans* interactions. (A, B) Representative steady-state patterns of the multicellular system from simulations with differentially labeled Ihog-expressing cells (A) and mixed Ihog- and Hh-expressing cells (B). (C) The average numbers of total neighbor cells and “like” neighbor cells were quantified for scenarios (A, I∼I bars) and (B, I∼H bars). Data were obtained from 300 random snapshots. Each error bar shows the mean ± SD. (D, E) Blue lines are the quantified relationships between the average numbers of total neighbor cells (D) and “like” neighbor cells (E) in a mixed system as a function of the difference in strength between the heterophilic Ihog-Hh (*γ_IH_*) and homophilic Ihog-Ihog (*γ_II_*) *trans* interactions. For comparison, the red dashed lines mark the values obtained from homogeneous system with differentially labeled Ihog-expressing cells. The shaded areas outline the standard deviations around the corresponding central average values. Each data point was calculated using 300 random snapshots from the simulation. (F) Simulation with *γ_IH_*: *γ_II_* = 30, starting with an aggregate formed from 50 Ihog-expressing cells, then 50 Hh-expressing cells were added into the simulation space (left). The energy-based evolution leads to the surface engagement of Hh-expressing cells onto the Ihog-expressing cell aggregate (middle) and eventually a checkerboard-like morphological pattern (right) appears as the simulation reaches steady state. (G) S2 cells were transfected with plasmids expressing Hh and GFP or Ihog and mCherry as indicated. 48 hours after transfection, cells were resuspended by pipetting and then mixed as indicated. Cell mixture was incubated with gentle rotating for 12 hours. At the indicated time points, an aliquot of cells was removed and imaged with a confocal microscope. Representative images from 3 experiments are shown. Scale bar, 100 µm.

We simulated 50 Ihog-expressing cells and 50 Hh-expressing cells (Fig. 7B). When we altered the ratio of the heterotypic and the homotypic interaction strength (*γ_II_*, *γ_IH_* values), the morphology of the mixed system changed. From these values and the patterns, we obtained the average number and type of neighbor cells for any given center cell. We found that when *γ_IH_* is 30 times larger than *γ_II_*, the mixed system exhibited the checkerboard-like morphological pattern (Fig. 6B), within which each center cell had 4.1 ± 0.7 neighbors, and only 0.3 ± 0.5 of them were “like” cells (Fig. 7C, I∼H bars).

This simulation study of the effect of the parameter *γ_II_*: *γ_IH_* predicted that the number and type of neighbor cells are sensitive to the *γ_IH_*: *γ_II_* ratio. A honeycomb-to-checkerboard transition occurred when *γ_IH_*: *γ_II_* is between 20 and 30 (Fig. 7D, E). Therefore, the neighbor statistics from the experimental aggregation assay and the computational modeling suggested that the relative affinity of the Ihog-Hh heterotypic interaction is at least ∼ 20 – 30 times higher than that of the Ihog-Ihog homotypic *trans* interaction. Previous sedimentation velocity and sedimentation equilibrium experiments predicted that the dimerization constants for the extracellular portion of Ihog comprising Fn1–2 range from 60 to 430 μM and that the dissociation constants for the interaction between HhN and this IhogFn1–2 region were between 0.4 – 8.0 μM (McLellan et al. 2006; Zhang et al. 2007).. Although we cannot differentiate homophilic *cis* interactions from homophilic *trans* interactions, our modeling prediction of an affinity for the Ihog-Hh heterotypic interaction that is at least 20 times higher than that for the Ihog-Ihog homotypic *trans* interaction is consistent with the values obtained in the biochemical experiments (McLellan et al. 2006; Zhang et al. 2007).

### Heterophilic Ihog-Hh binding displaces pre-established homophilic Ihog *trans* interactions

Differential adhesion has been proposed to promote cell rearrangements in cell aggregates such that strongly adhesive cells sort together and weakly adhesive cells are excluded (Foty and Steinberg 2005; Steinberg 2007). Thus, from our computational analysis of interaction strength, we predicted that mixing Hh-expressing cells with pre-existing aggregates of Ihog-expressing cells would displace the relatively weaker Ihog-mediated homophilic interactions by establishing stronger Hh-Ihog interactions and result in a checkerboard pattern. To test this, we performed both computational model simulations and experiments with S2 cells. With *γ_IH_*: *γ_II_*, =30 in the computational model, we observed the development of the checkerboard pattern as the simulation reached steady state (Fig. 7F).

In the S2 cell experiment, we mixed differentially labeled Hh-expressing cells with preformed aggregates of Ihog-expressing cells (Fig. 7G). Within 30 minutes after cell mixing, Hh-expressing cells were found on the surface of the pre-existing aggregates of Ihog-expressing cells, which we interpreted as Hh binding to Ihog proteins that were not engaged in homophilic adhesion. Over time, Hh-expressing cells were found inside the Ihog-expressing cell aggregates, such that, by 12 hours after mixing, all cell aggregates contained similar numbers of Hh-expressing cells and Ihog-expressing cells arranged in a checkerboard-like pattern (Fig. 7G). This pattern is consistent with cell rearrangements caused by differential adhesion with the Hh-Ihog interaction exhibiting a higher affinity than the Ihog-Ihog homophilic *trans* interaction. Moreover, the observed cellular rearrangement indicated the Hh ligand disrupts *trans* Ihog-Ihog binding by competing for the Ihog Fn1 domain. Taken together, these results suggested Hh binding to Ihog is dominant over Ihog-Ihog homophilic interactions and effectively competes for Ihog even in the context of pre-established Ihog-Ihog *trans*-homophilic interactions.

## DISCUSSION

We investigated the functional roles of the Hh receptor Ihog by determining a mechanism by which the Ihog proteins stabilizes cytonemes in the *Drosophila* wing imaginal disc. We found a dual role for cytoneme-localized Ihog proteins in Hh signal transduction and in *trans*-homophilic binding that mediates cytoneme-cytoneme interactions. This dual function is not unique to the Ihog proteins. Like Ihog, other members of the Ig-CAM family, such as the netrin receptor DCC, the Slit receptor Robo, and neural cell adhesion molecule N-CAM, also have dual roles in adhesion and signaling (Juliano 2002; Orian-Rousseau and Ponta 2008). The ligand binding and cell adhesion functions often involve different extracellular domains of the same protein, thus are biochemically separated and mutually compatible (Frei et al. 1992; Martin-Bermudo and Brown 1999; Sjostrand et al. 2007). However, the Ihog homophilic binding site overlaps with Ihog-Hh interface on Fn1 of the Ihog extracellular domain, resulting in competition between Hh binding and Ihog-Ihog interactions.

On the basis of these two functions competing for an overlapping surface on the Ihog Fn1 domain and the differential interaction strength between Ihog-Hh heterophilic and Ihog-Ihog *trans*-homophilic bindings, we propose a model in which Ihog-Ihog *trans* interactions promote and stabilize direct cytoneme-cytoneme contacts to facilitate Ihog in reaching and capturing the Hh ligands secreted from the cytonemes. The stronger ligand-receptor interaction releases Ihog from the weaker *trans*-homophilic interaction, enabling the receptor-ligand complex to freely transport or become internalized along the cytonemes. Meanwhile, the unbound former homophilic binding-partner Ihog either forms a new homophilic contact or engages in ligand-receptor complex formation along the cytonemes (Fig. 8). Thus, the apparently incompatible functions of Ihog in homophilic adhesion and ligand binding cooperate to promote Hh transport and reception along the cytonemes. The model incorporates the role of the glypicans, Dlp and Dally. Although contrary to a previous model that limited the contribution of the glypicans in cytoneme stabilization to a trans role (Gonzalez-Mendez et al., 2017), in our model Dlp or Dally can contribute either as the membrane-tethered or the shed form and participate Ihog-Ihog trans interactions. Our model is also consistent with the reported affinity of Hh for the Ihog-Ptc receptor complex, which is higher than the affinity of Hh for the co-receptor Ihog alone (Zheng et al. 2010). Indeed, the presence of Ptc in the Hh-receiving cytoneme is critical to Hh reception in the responding cells (Chen et al. 2017). Thus, we propose that the integration of the functions of Ihog — promotion of cytoneme-cytoneme contacts, Hh delivery, and Hh signal reception — depends on the differential affinity (Ihog-Ihog < Ihog-Hh < Ptc-Ihog-Hh) and the competitive binding between Ihog for itself (in *trans*) and Hh (Fig. 8).

**Figure 8.**
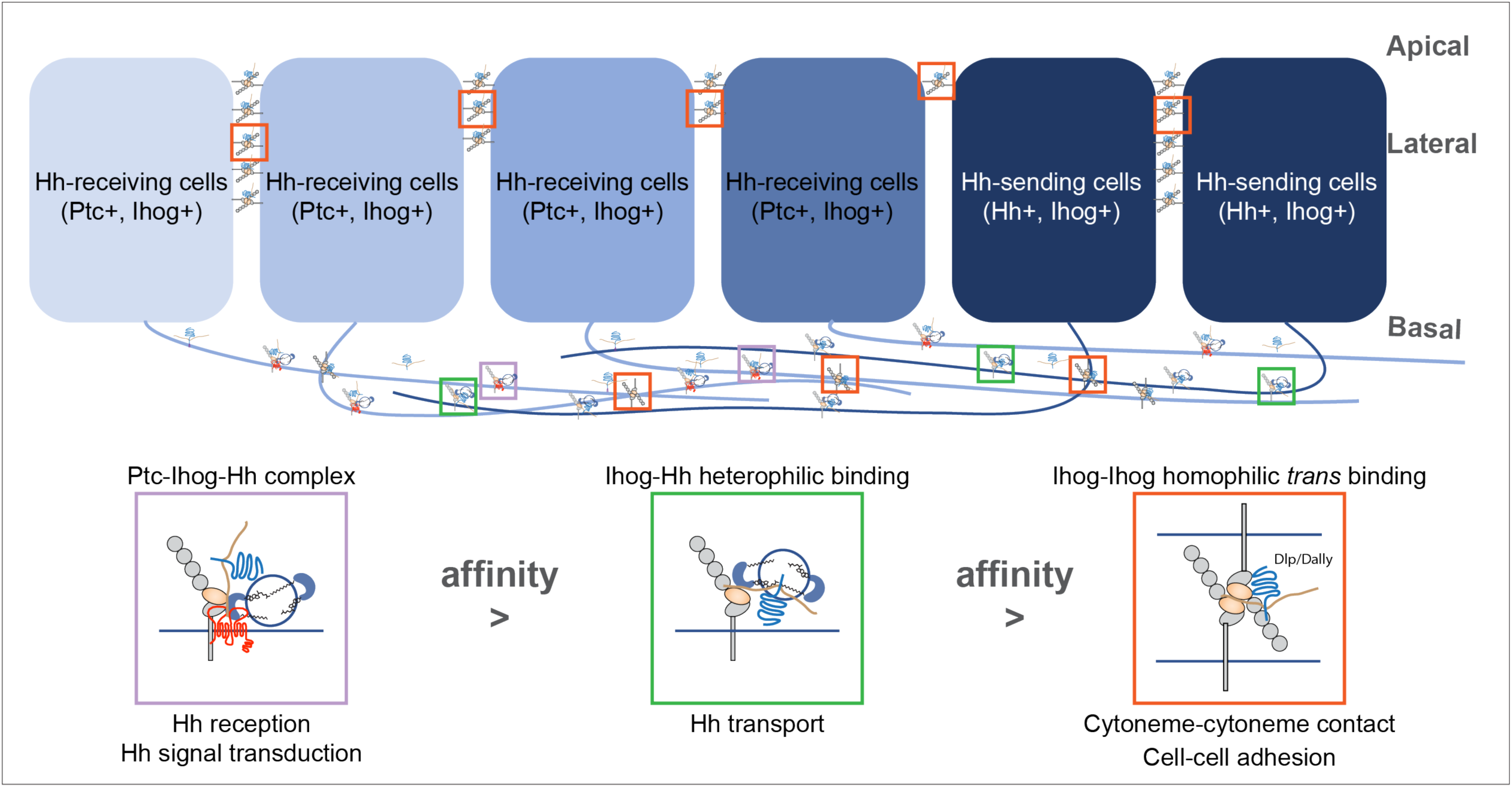
A model of the competitive coordination of the dual roles of Ihog in homophilic adhesion and signal reception. Diagram illustrating Ihog-Ihog homophilic *trans* interaction and Ihog-Hh heterophilic interaction in the wing imaginal disc epithelium. For simplicity, only a small number of the apical and lateral interactions are shown. Based on the differential affinity (Ptc-Ihog-Hh > Ihog-Hh > Ihog-Ihog) and the competitive binding between Ihog for itself (in *trans*) and for Hh, we propose a model in which Ihog-Ihog *trans* interactions promote and stabilize cytoneme-cytoneme contacts, thereby facilitating the “capture” of Hh ligands released by cytonemes of Hh-expressing cells by Ihog on cytonemes of adjacent cells, ultimately reaching Ihog on cytonemes of Hh-receiving cells. The stronger Hh-Ihog interaction triggers release of Ihog from the weaker *trans*-homophilic interaction, enabling the receptor-ligand complex to transport along the cytoneme. Ultimately, the strongest interaction of Hh with the Ptc and Ihog complex results in Hh signal transduction. Both Hh release and cytoneme formation occur at the basal side of the wing disc epithelium, making weaker Ihog-Ihog *trans* interactions accessible for the replacement by stronger Ihog-Hh interactions along the cytonemes, where Ihog also functions as the receptor for Hh transport and reception. In contrast, farther from the source of secreted Hh, heterophilic Ihog-Hh interactions would be infrequent along the lateral side of epithelia, where the *trans* Ihog-Ihog interaction plays an essential role in maintaining A/P cell segregation and lineage restriction. The heparan sulfate necessary for the Ihog-Hh or *trans* Ihog-Ihog interactions may be supplied by Dally or Dlp, either as the membrane-associated forms of these glypicans or as the form released upon shedding.

The *trans*-homophilic Ihog-Ihog interaction is not only critical for cytoneme stabilization but also for A/P compartment boundary maintenance in the *Drosophila* wing discs (Hsia et al. 2017). Notably, both Hh release and cytoneme formation occurs at the basal side of the wing disc epithelium (Callejo et al. 2011; Bilioni et al. 2013; Bischoff et al. 2013; Gradilla et al. 2014; Chen et al. 2017; Gonzalez-Mendez et al. 2017), making weaker Ihog-Ihog *trans* interactions accessible for replacement by stronger Ihog-Hh interactions along the cytonemes, where Ihog also functions as the receptor for Hh transport and reception (Fig. 8). In contrast, farther from the source of secreted Hh, heterophilic Ihog-Hh interactions would be infrequent along the lateral side of epithelia, where the *trans* Ihog-Ihog interactions play essential roles in modulating A/P cell segregation and lineage restriction (Hsia et al. 2017). In addition, direct membrane contacts are much more extensive along the lateral sides of epithelia compared with that formed along cytonemes (Fig. 5), favoring persistent Ihog-Ihog *trans* interactions that create an additional barrier for direct competition from the basally released Hh ligands. Thus, in agreement with the functional needs and the availability of Hh ligands, Ihog-Ihog homophilic *trans* interactions along the cytonemes are dynamic and readily switchable, whereas those at the lateral cell-cell junctions are more stable and less likely to be disrupted (Fig. 8).

Beside the wing imaginal disc epithelia, cytoneme-mediated Hh reception and transport has been described in other *Drosophila* tissues, such as the abdominal and the female germline stem cell niche (Rojas-Rios et al. 2012; Bischoff et al. 2013; Gonzalez-Mendez et al. 2017). The involvement of cytonemes in Hh signaling has been extended from insect to vertebrates by studies of the limb bud of chick embryo and cultured mouse embryonic fibroblasts (Sanders et al. 2013; Hall et al. 2020). The vertebrate homologs of the *Drosophila* Ihog proteins, Cdo and Boc, localize in cytonemes, and overexpression of either CDO or BOC increases the number of cytonemes detected on mammalian cells (Hall et al. 2020). Furthermore, cytonemes have been implicated in the delivery of other paracrine signaling molecules important in development, including Notch, epidermal growth factor, fibroblast growth factor, bone morphogenetic protein, and Wnt (Gonzalez-Mendez et al. 2019). Remarkably, cytonemes from a single cell often exhibit different receptor compositions such that different cytonemes from the same cell can selectively respond to ligands for a specific pathway and not others (Roy et al. 2011). The mechanism that segregates receptors to different cytonemes is yet not known. Whether homotypic adhesion contributes to the distinct localization of morphogen receptors to different cytoneme remains an open question. Further studies are necessary to explore whether the vertebrate Hh co-receptors CDO and BOC also have dual roles in adhesion and signaling and whether homophilic interactions mediated by this family of proteins contributes to additional signaling events.

## MATERIALS AND METHODS

### Antibodies

Antibodies and dilutions used were mouse anti-Dally-like (Dlp) antibody 1:50 (DSHB, 13G8); mouse anti-Ptc antibody 1:50 (DSHB, Apa1); mouse anti-*α*-tubulin (DM1A, EMD Millipore) 1:5000; mouse anti-HA 1:1000 (HA.11, Covance); mouse anti-*β*-tubulin 1:5000 (DSHB, E7); rabbit anti-GFP 1:1000 (Invitrogen, A-11122); rat anti-Ihog antibody 1:500 (Yao et al. 2006); rabbit anti-Hh at 1:500 (Tabata and Kornberg 1994). HRP-conjugated and Fluorophore-conjugated secondary antibodies were from Jackson Immuno-Research Lab. Alexa Fluor® 594 Phalloidin was from Thermo Fischer. The antibody information is also listed in the key resources table (Supplementary Table 1).

### Cell culture and transfection

*Drosophila* S2 cells (DGRC) were cultured in *Drosophila* Schneider’s medium supplemented with 10% of fetal bovine serum (Omega Scientific) and 1% Penicillin-Streptomycin-Glutamine (Thermo Fisher) at 25°C in a humidified incubator. Transfection was performed with FuGENE 6 transfection reagent (Promega). Expression constructs of GFP, mCherry, HhN, Hh, Ihog and Ihog-YFP used in *Drosophila* cell culture were cloned into pAcSV plasmid as previously described (Wu et al. 2019).

### Cell aggregation assay

S2 cells were transfected separately with plasmids expressing desired proteins. 48 hours after transfection, S2 cells were washed with PBS and dissociated by 0.05% trypsin treatment for 5 min at 25°C. The dissociated cells were resuspended in fresh medium with 10% fetal bovine serum. The resuspended cells were then incubated in 1.5 ml ultra-low adhesion microcentrifuge tubes with gentle rotation at room temperature for the time indicated in the figure legends. Cells were then transferred into glass bottom dishes (D35-20-1.5-N, In Vitro Scientific) for live imaging by microscopy. In the experiments involving mixing differentially labeled red and green cells, cells co-expressing GFP or mCherry with the plasmid expressing the protein of interest were counted under microscope and mixed with equal number of transfected cells prior to incubation with rotation.

To assess cell aggregation, low-magnification fields of similar cell density were randomly taken from each cell aggregation experiment, and the cell clusters were scored if they contained three or more cells. The aggregation effect was quantified as the ratio of certain transfected cells within clusters to total transfected cells (both clustered and non-clustered). Each bar shows the mean ± SD from 20-30 different images. Unpaired two-tailed t test was used for statistical analysis. Statistical analysis was performed using GraphPad Prism software.

### Cell immunostaining and imaging

48 hours after transfection, dissociated S2 cells were allowed to settle and adhere for 60 min on a glass coverslip. Cells were then washed twice with PBS, fixed in 4% formaldehyde (Electron Microscopy Sciences) in PBS, blocked and permeabilized by 1.5% normal goat serum (NGS) & 0.1% Triton X-100 in PBS, incubated with primary antibody in PBS containing 1.5% NGS and 0.1% Triton X-100 for 1 hr at room temperature, washed 3 times with 0.1% Triton X-100/PBS, incubated with secondary antibody (with or without Phalloidin) and washed with 0.1% Triton X-100/PBS. The stained cells were mounted using the Vectashield anti-fade mounting medium (H-1000) and imaged with a Zeiss spinning disc confocal microscope.

### Computational model of cytonemes

We simulated the cytonemes using a *in silico* stochastic assay. The cell surface is simplified as a linear base line with length *M* (shown as the black solid lines in the bottom of Fig. 5A). In the initial step, we randomly picked 30% × *M* locations along the base line as the initial locations for the cytonemes. The initial cytoneme lengths are all 0.

In each simulation step, the binary interaction variable 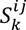 was randomly assigned for each pair of neighboring segments according to Eq. 3. From 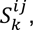, the number of tip interactions for each cytoneme (*T_i_*) and the total homophilic *trans* interaction for the *i^th^* cytoneme 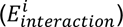 can be obtained. For each cytoneme, its probabilities of elongation and shrinkage (i.e. *p_elongation_* and *p_shrinkage_*) were computed using equations 1 & 2; a random number *r* ∈ [0, 1] was picked; if the cytoneme length was larger than 0 and *r* ≤ *p_shrinkage_* / *p_elongation_* + *p_shrinkage)_*, the cytoneme length was shrunk by 1 segment; otherwise it was elongated by 1. Once all cytonemes were elongated/shrunk, 1 cytoneme was randomly picked and tentatively moved along the cell surface base line; the interaction variables 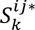 were re-assigned based this tentative configuration and the energy change 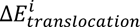 was calculated; another random number *r* ∈ [0, 1] was picked and the tentative move would be accepted if and only if 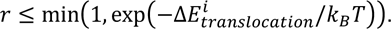.

For each choice of *E_ii_*, we allow the cytoneme system to evolve for more than 5,000,000 steps. Data were collected after 1,000,000 simulation steps when the cytoneme system reached steady state. In our showed results, we set 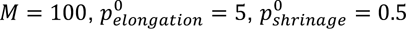 and *α* = 0.2. We tested different parameters and the general conclusions remained the same.

The cytoneme bundles were identified as a collection of parallel cytonemes located close together and forming more than 3 pairwise contacts between two cytonemes. The cytoneme bundling index is calculated by multiplying the minimum cytoneme length by the cytoneme number within each bundle.

### Computational modeling of cell rearrangement

We modeled the Ihog-Ihog and Ihog-Hh interactions using a vertex-based *in silico* assay (Bi et al. 2015; Park et al. 2016). Based on mechanical free energy calculated according to Eq. 5, we used the Metropolis Monte Carlo method to perform the simulations. Simulations were performed within a *L* × *L* 2-D square space with a periodic boundary condition along the x- and y-axes. The initial conditions were sets of randomly generated morphologies: We first assigned *N* cellular points and 5 × *N* environmental points randomly distributed in the 2-D space, then used the Voronoi tessellation function in MatLab to partition the space into 6 × *N* polygons based on these points, representing the *N* cells and the empty space surrounding them.

We implemented Metropolis Monte Carlo simulations by moving the 6 × *N* points according to the mechanical energy: In each tentative move, a random point is selected and a random displacement *Δ**l*** is assigned to it; the moved point causes re-partitioning of the 2-D monolayer using the Voronoi tessellation function, which results in a change in the monolayer’s mechanical energy, *ΔU*’; a random number *r* ∈ [0, 1] is picked and the tentative move will be accepted if and only if *r* ≤ min(1, exp(−*ΔU*’/*k_B_T*)).

We allowed the monolayer to randomly evolve for more than 650,000 steps. To ensure simulation efficiency, we adjusted the maximum allowed displacement to maintain the overall accept rate to be around 25∼40%. Data were collected after 200,000 simulation steps when the monolayer’s morphology had reached steady state.

In our simulations, *L* is set to 20, and both the Hh-expressing S2 cells and the Ihog-expressing S2 cells are set with a unit area *A*_0_ = 1. For simplicity, we assumed that areal elastic coefficient *α* = 500 and the contractile coefficient *β* = 6 are the same for different types of cells. To account for the differential cell-cell adhesion, *γ_HH_* was set to 0 (Hh-expressing cells do not aggregate), *γ*_*II*_ = 0.25 for interaction between Ihog-expressing cells, we modulated the ratio *γ_IH_*: *γ_II_* to set *γ_IH_* for our parameter study.

### Drosophila strains

The *ptc-GAL4 (Hinz et al. 1994)* driver and *tub-Gal80^ts^ (McGuire et al. 2003)* were used for transient expression of transgenic constructs using the GAL4/UAS system (Brand and Perrimon 1993). Fly crosses were maintained at 18°C, and the *Gal80^ts^* repressor was inactivated for 24 hr at restrictive temperature (29°C) before dissection. The *actin>y+>GAL4* (Ito et al. 1997) driver was used to generate random ectopic clones of the *UAS* lines. The *CoinFLP-LexA::GAD.GAL4* driver was used to generate random clones expressing either *UAS-CD4-spGFP1-10* or *lexAop-CD4-spGFP11* (Feinberg et al. 2008; Gordon and Scott 2009; Bosch et al. 2015). Larvae of the corresponding genotypes were incubated at 37°C for 30-60 min to induce *hs-FLP*-mediated recombinant clones. The genotypes (see **DETAILED GENOTYPES**) of larvae for transient or random expression of transgenic constructs are listed in the key resources table (Supplementary Table 1).

### Imaginal discs immunostaining and imaging

Wing discs from 3^rd^ instar larvae were dissected, fixed in 4% formaldehyde (Electron Microscopy Sciences) in PBS, blocked and permeabilized by 5% normal goat serum (NGS) & 0.3% Triton X-100 in PBS, incubated with primary antibody in PBS containing 5% NGS and 0.3% Triton X-100 overnight at 4°C, washed 3 times with 0.3% Triton X-100/PBS, incubated with secondary antibody, and washed with 0.3% Triton X-100/PBS. The stained discs were mounted and imaged with a ZEISS spinning disc confocal microscope or a ZEISS LSM 880 with Airyscan. Average cytoneme length was determined using ImageJ and plotted using GraphPad Prism software.

### MBP-HhN purification

The MBP-HhN expression plasmid was a gift from Dr. Daniel Leahy (The University of Texas at Austin). A DNA fragment encoding the *Drosophila melanogaster* Hh residues 85–248 (HhN) was cloned into the MBP-HTSHP expression vector, which was modified based on the pMal-c2x vector (New England Biolabs) by including a linker region with various tags (His-TEV-Strep-His-PreScission). Similar to the procedure described previously (8), the fusion proteins were expressed in *Escherichia coli* strain B834 (DE3) by induction with 1 mM isopropyl 1-thio-*β*-D-galactopyranoside overnight at 16 °C. Cells were harvested, lysed, and centrifuged, and the supernatant was passed over nickel-nitrilotriacetic acid resin (Qiagen). Proteins were eluted with imidazole according to the manufacturer’s suggestions. The elution was then placed into 6000 – 8000 molecular weight–cutoff 40-mm dialysis tubing and dialyzed against 20 mM Tris (pH 8.0) and 200 mM NaCl.

### Western blot analysis

48 hours after transfection, S2 cells were lysed in 1% NP40 (50 mM Tris-HCl at pH 6.8, 150 mM NaCl, and protease inhibitors) for 30 min at room temperature. The lysate was clarified by centrifugation, and proteins were recovered directly in SDS-PAGE sample buffer. Proteins were separated by SDS-PAGE under reducing conditions and then transferred onto PVDF membranes (Millipore). After protein transfer, the membranes were blocked and then immunostained with primary antibodies and HRP-conjugated secondary antibodies.

### Statistical analysis

All data in column graphs are shown as mean values with SD and plotted using GraphPad Prism software. Statistical analyses were performed with unpaired two-tailed t-test, one-way ANOVA followed by Dunnett’s, Sidak’s or Tukey’s multiple comparisons test, or two-sided Fisher’s exact test was used for statistical analysis as described in the figure legends. The sample sizes were set based on the variability of each assay and are listed in the Figure legends. Independent experiments were performed as indicated to guarantee reproducibility of findings. Differences were considered statistically significant when P < 0.01.

## ACKNOWLEDGMENTS

We thank S. Blair, I. Guerrero, D. Leahy, T. Tabata, the Bloomington *Drosophila* Stock Center, the Vienna *Drosophila* RNAi Center, Kyoto Stock Center and the Developmental Studies Hybridoma Bank for fly strains and reagents. We thank A. Popratiloff from the GW Nanofabrication & Imaging Center, members of the Anatomy and Cell Biology Department for comments during the development of this work. We thank N. R. Gough for helpful discussions.

## AUTHOR CONTRIBUTIONS

X.Z. and G.L. conceived of the presented idea. S.Y., Y.Z, X. W., X.S.W., R.C and X.C. carried out the experiments. G.L., C.Y, and S.O. developed the theory and performed the computations. All authors discussed the results and contributed to the final manuscript.

---

**Supplementary Figure 1.**
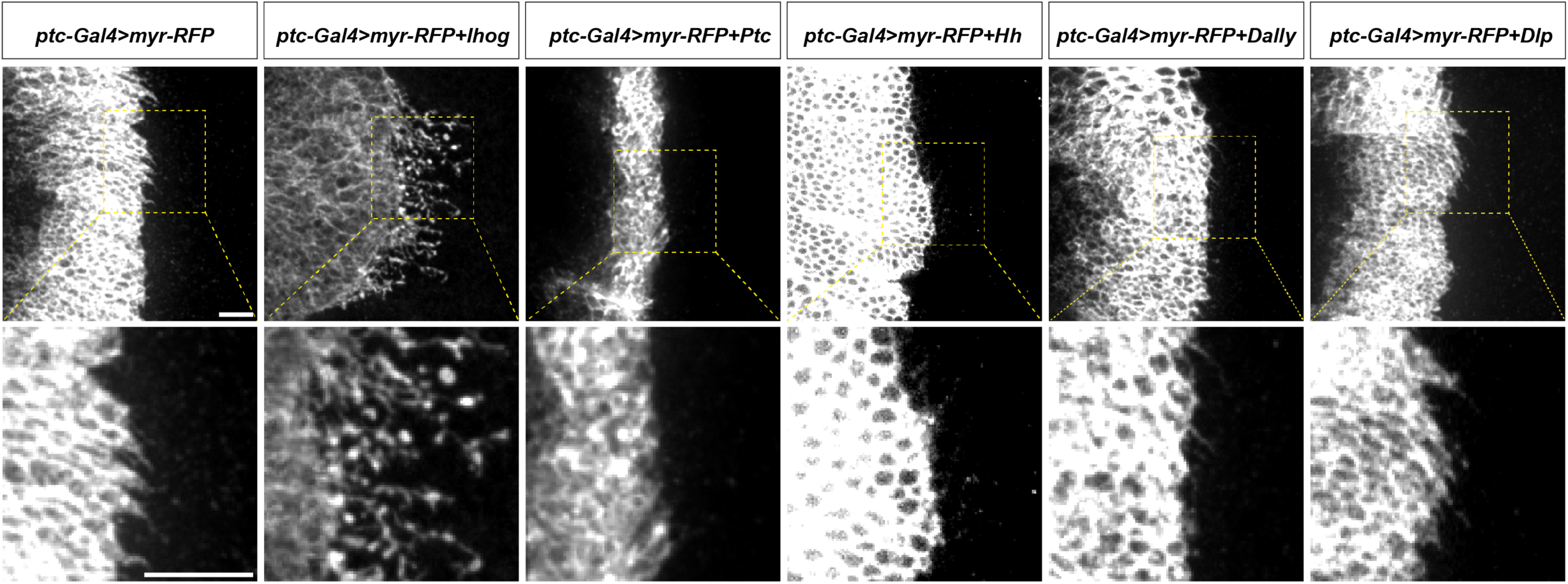
Ectopic expression of Ihog, but not other Hh pathway components, stabilizes cytonemes. Wing discs from 3^rd^ instar larvae carrying *ptc-GAL4, tub-GAL80^ts^* and the indicated *UAS-*transgenes were immunostained for RFP to visualize cytonemes projecting from Hh-receiving cells. *UAS-Myr-RFP*: myristoylated form of red fluorescent protein that marks the cell membrane and enables visualization of the cytonemes. Scale bar, 10 µm.

**Supplementary Figure 2.**
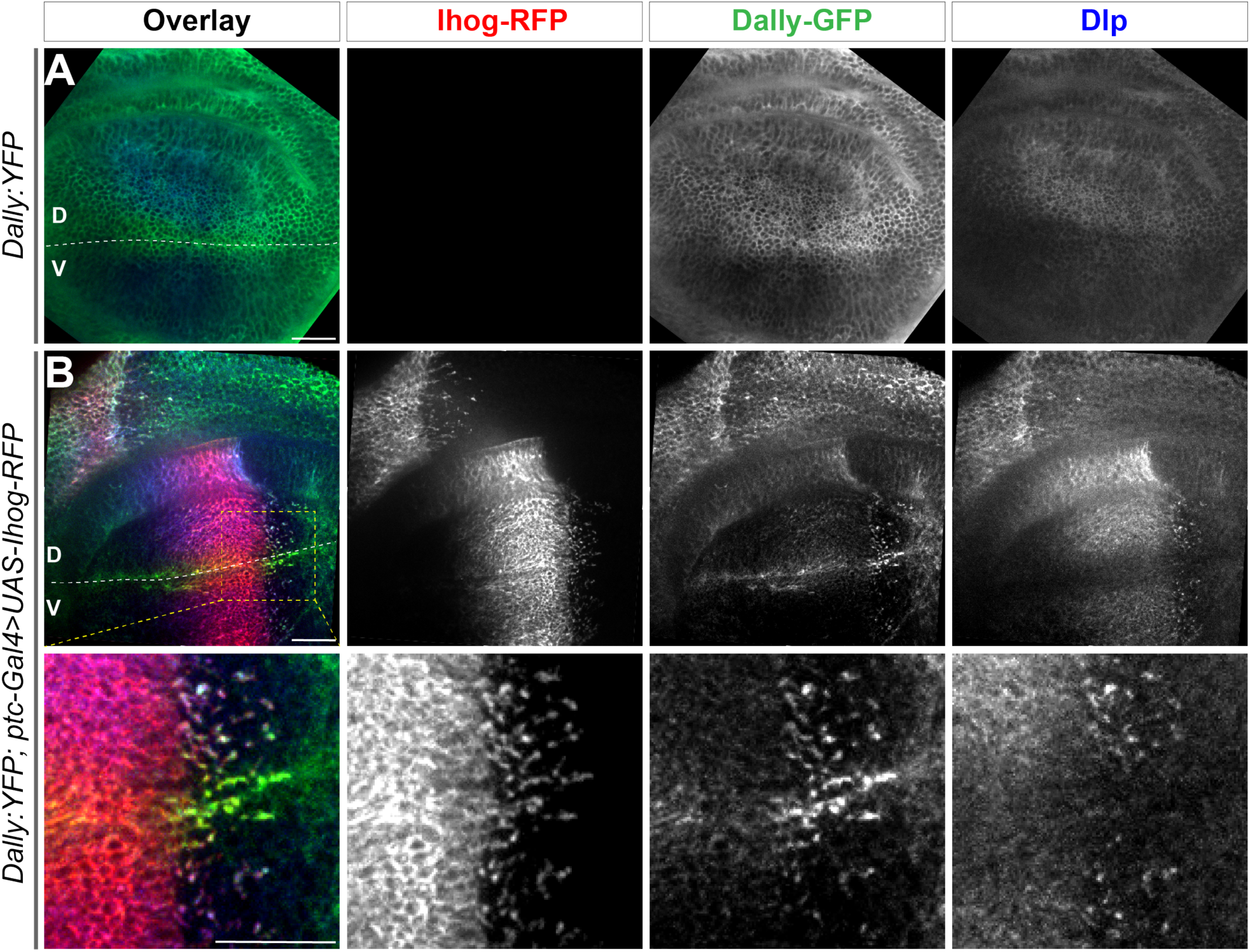
Ectopic Ihog induces accumulation of Dally and Dlp that is associated with the different distributions of the two glypicans. ((A) Distribution of Dally:YFP and Dlp in a wing discs from a 3^rd^ instar larvae carrying a Dally:YFP protein trap allele. (B) Distribution of Dally:YFP, Dlp, and Ihog in a wing disc from larvae expressing the a Dally:YFP protein trap allele and the *ptc-GAL4, tub-GAL80^ts^* and *UAS-Ihog-RFP* transgenes. The lower row shows an enlarged area (outlined in upper left panel) around the D/V boundary. The discs were immunostained to visualize Dlp (blue) and with antibody against YFP to visualize Dally (green). Ihog-RFP was visualized with the RFP fluorescence (red). The dorsal/ventral (D/V) boundary is indicated by a dashed white line. Scale bar, 20 µm.

**Supplementary Figure 3.**
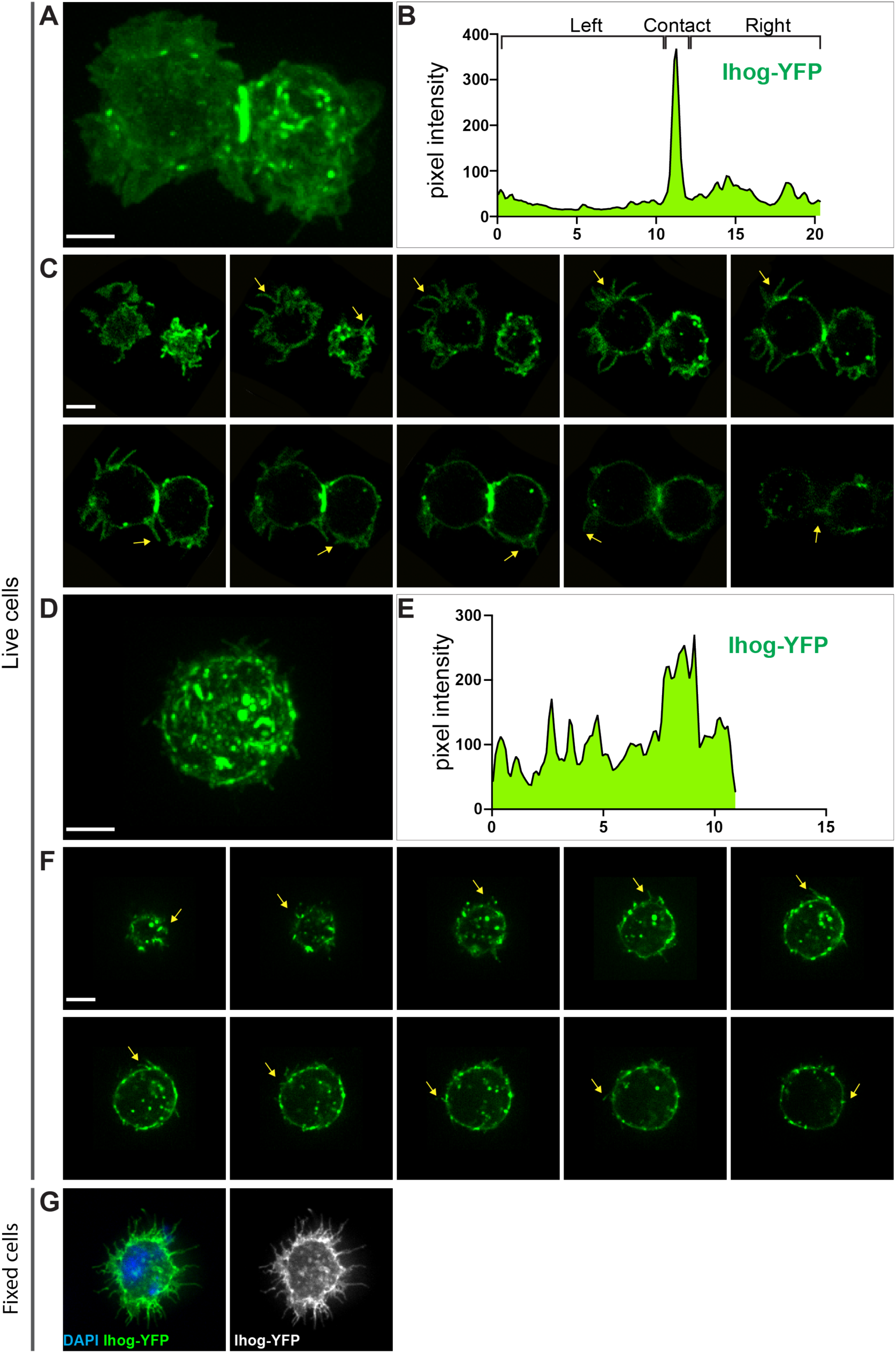
Subcellular localization of Ihog proteins in live S2 cells. (A-F) Live imaging of S2 cells transfected with plasmids expressing YFP-tagged Ihog was performed and pixel intensity plotted. Images of xy focal planes were collected along the z-axis through the S2 cells with each image consisting of a maximum projection of three-dimensional volume encompassing the entire cell(s). (A, B) Representative maximum intensity projections of z-stack sections of S2 cells in contact and pixel intensity of Ihog-YFP across the image. Data are representative of n > 3 experiments. (C) Images from 10 different xy focal planes with identical z-axis distance for the cells shown in C. Arrows indicate filopodia. See Supplementary Movies 1, 2. (D, E) Representative maximum intensity projections of z-stack sections of single S2 cells and pixel intensity of Ihog-YFP across the image. Data are representative of n > 3 experiments. (F) Images from 10 different xy focal planes with identical z-axis distance for the cell shown in D. See Supplementary Movies 3, 4. (G) S2 cells transfected with plasmids expressing Ihog-YFP were plated on the glass surface, and stained for YFP following MEM-fix as previously described (73). Scale bar, 5 µm.

**Supplementary Figure 4.**
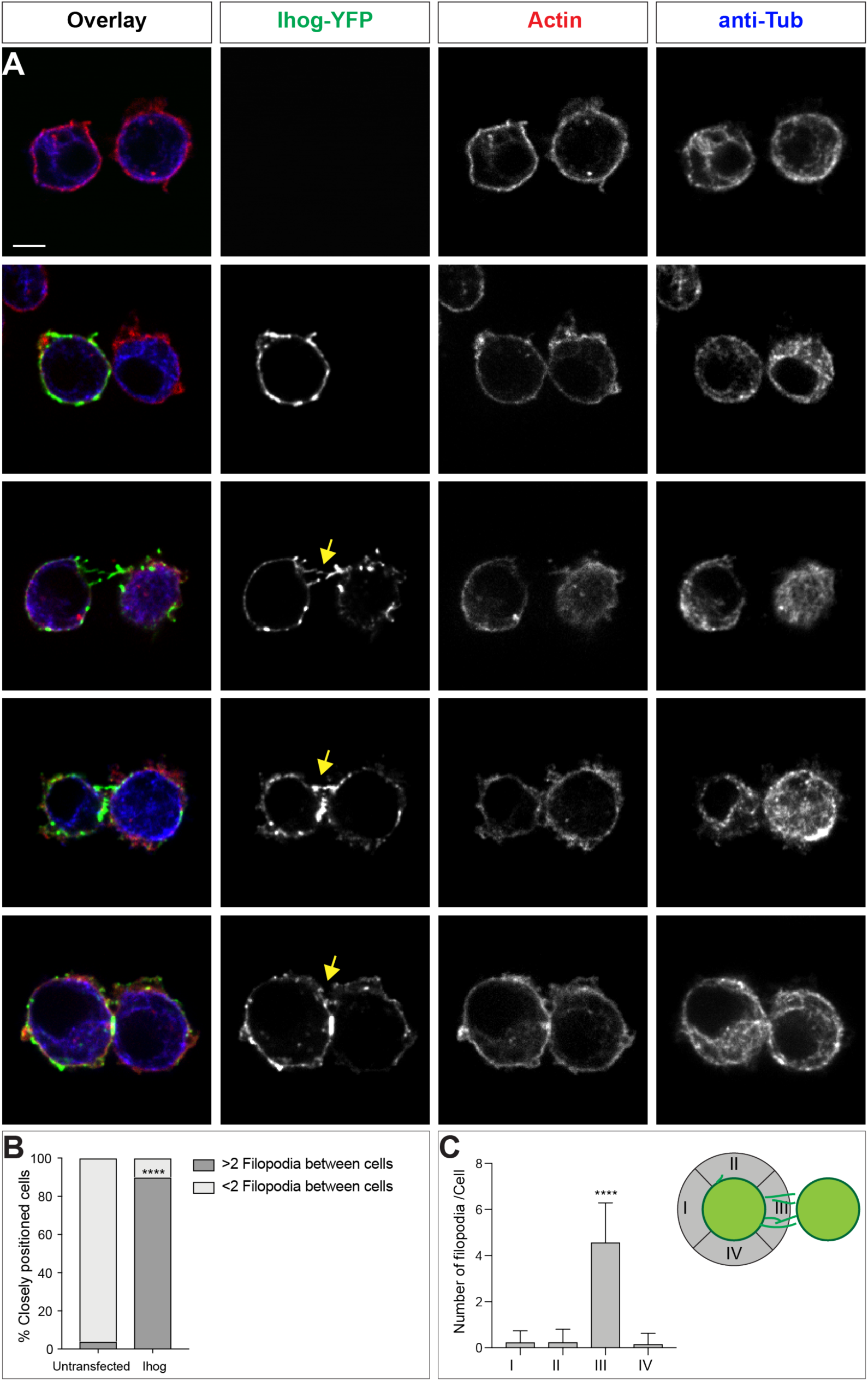
Ihog is enriched in filopodia-like structures of closely positioned Ihog-expressing cells. S2 cells transfected with plasmids expressing Ihog-YFP were plated on the glass surface, fixed, and stained for actin (Phalloidin, red) and alpha-tubulin (blue). Representative images of two closely positioned untransfected cells (A), singular Ihog-expressing cell located next to an untransfected cell, two closely positioned Ihog-expressing cells (C-D), and Ihog-expressing cells that appear to start forming a stable contact (E). Scale bar, 5 µm. (F) Quantification of the presence and absence of filopodia between closely located S2 cells with or without Ihog expression. Two-sided Fisher’s exact test was used for statistical analysis. ****P < 0.0001. (G) Quantification of filopodia spatial distribution of Ihog-expressing cells closely located to another Ihog-expressing cell. one-way ANOVA followed by Tukey’s multiple comparison test was used for statistical analysis. ****P < 0.0001.

**Supplementary Figure 5.**
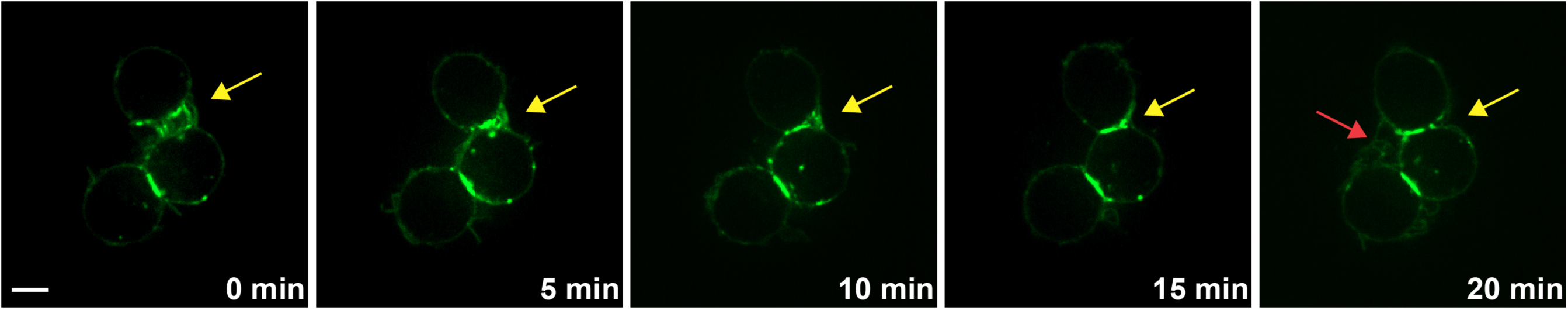
Live imaging shows that cell-cell contact occurs at sites of interaction between Ihog-enriched filopodia. Time-lapse imaging of S2 cells expressing Ihog-YFP. Interdigitation of filopodia at the region where Ihog-expressing cells initiate contact formation (yellow arrow). Progressively, filopodia are replaced by stable extensive contact between opposing cell membranes. Red arrow indicates a new cell-cell contact initiated by Ihog-YFP-positive filopodia. Scale bar, 5 µm.

**Supplementary Figure 6.**
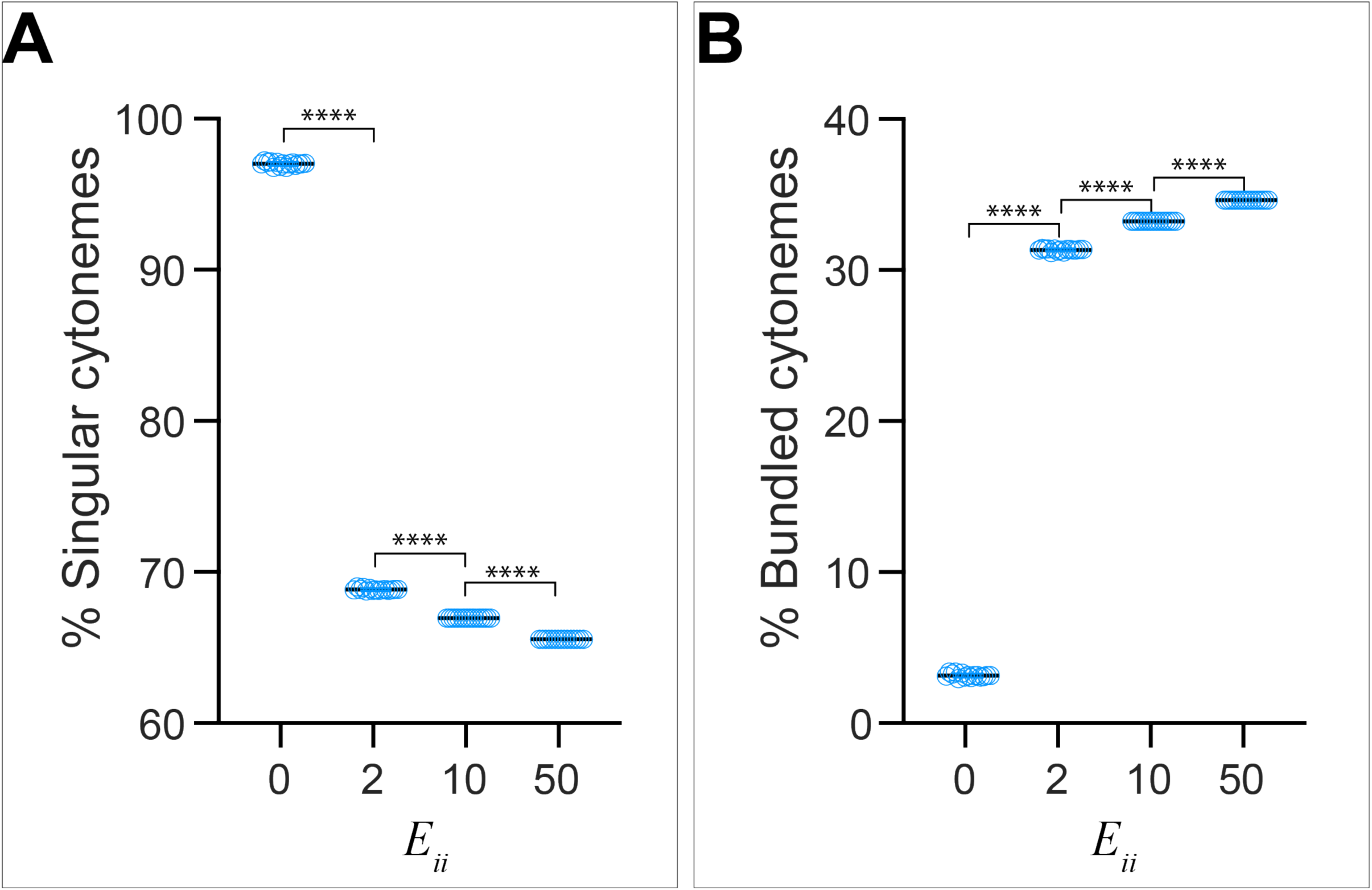
Effect of homophilic *trans*-interaction strength on the frequency of cytoneme bundle formation. The ratio of singular cytonemes (A) and the ratio of cytonemes present in bundles (B) from n=1001 random snapshots, each containing 30 cytonemes, are plotted against *E_ii_* ranging from 0 to 50. Each bar shows the mean ± SD, one-way ANOVA followed by Sidak’s multiple comparison test was used for statistical analysis. ****P < 0.0001.

**Supplementary Figure 7.**
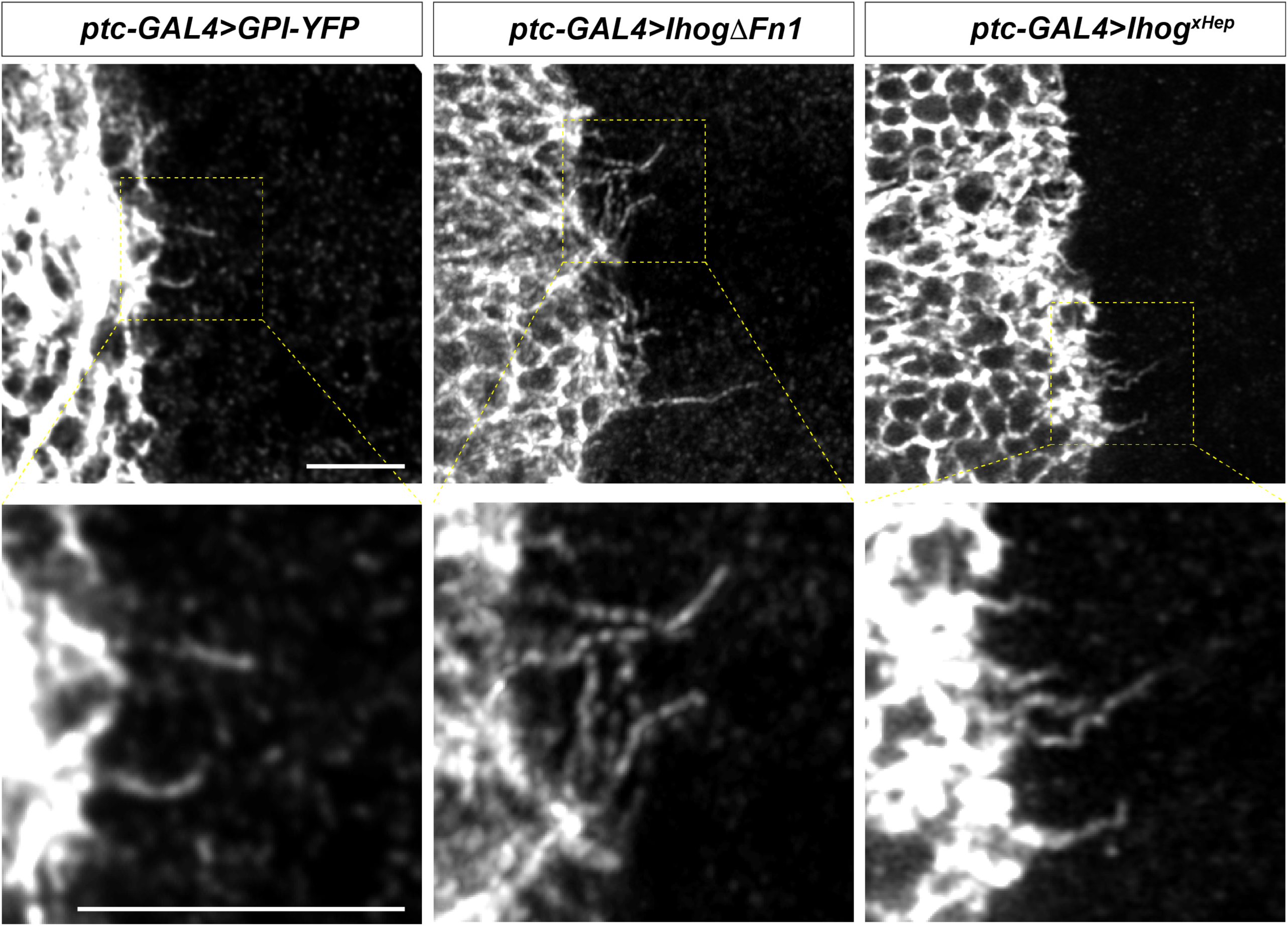
The Ihog Fn1 domain is essential for cytoneme bundling. Wing discs from 3^rd^ instar larvae carrying *ptc-GAL4, tub-GAL80^ts^* and the indicated *UAS-transgene* were immunostained and imaged with Airyscan. Scale bar, 5 µm.

**Supplementary Figure 8.**
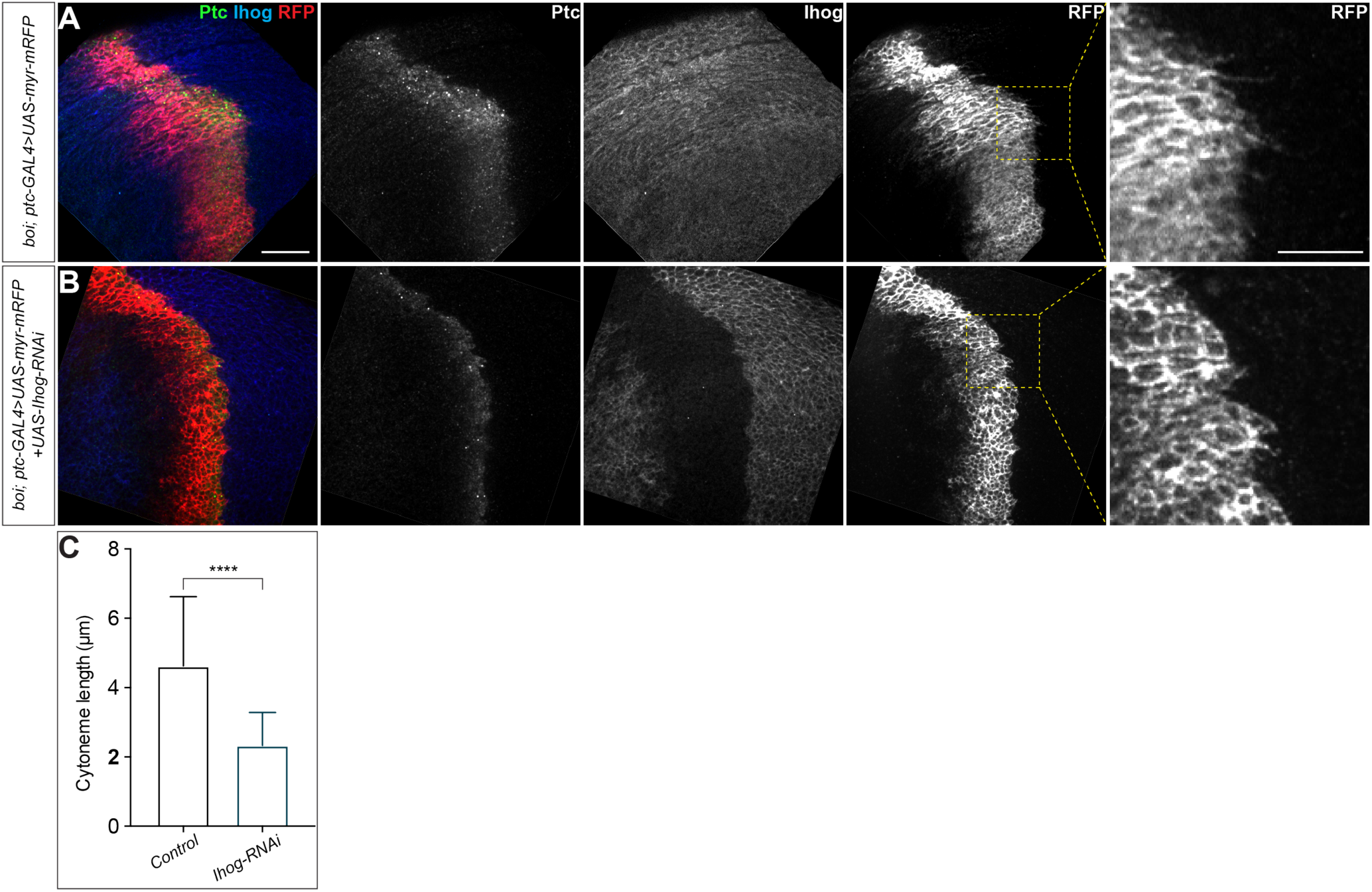
Loss of *Ihog* and its close paralogue *boi* from the wing imaginal discs cells resulted in cytonemes with reduced length. (A, B) Wing discs from 3^rd^ instar *boi* mutant larvae carrying the *ptc-GAL4, tub-GAL80^ts^* and the indicated UAS transgenes were immunostained to visualize Ptc (anti-Ptc, green), Ihog (anti-Ihog, blue), and the cytonemes (anti-RFP, red). Note that expression of *UAS-Ihog-RNAi* lead to loss of Ihog expression in the Ptc expressing domain (B) and resulted in cytonemes with reduced length. Scale bar, 20 µm. (C) Quantification of the average cytoneme length. Each bar shows the mean ± SD, n=30, and representative images are shown in (A, B). Two-tailed unpaired t-test was used for statistical analysis. ****P < 0.0001.

**Supplementary Figure 9.**
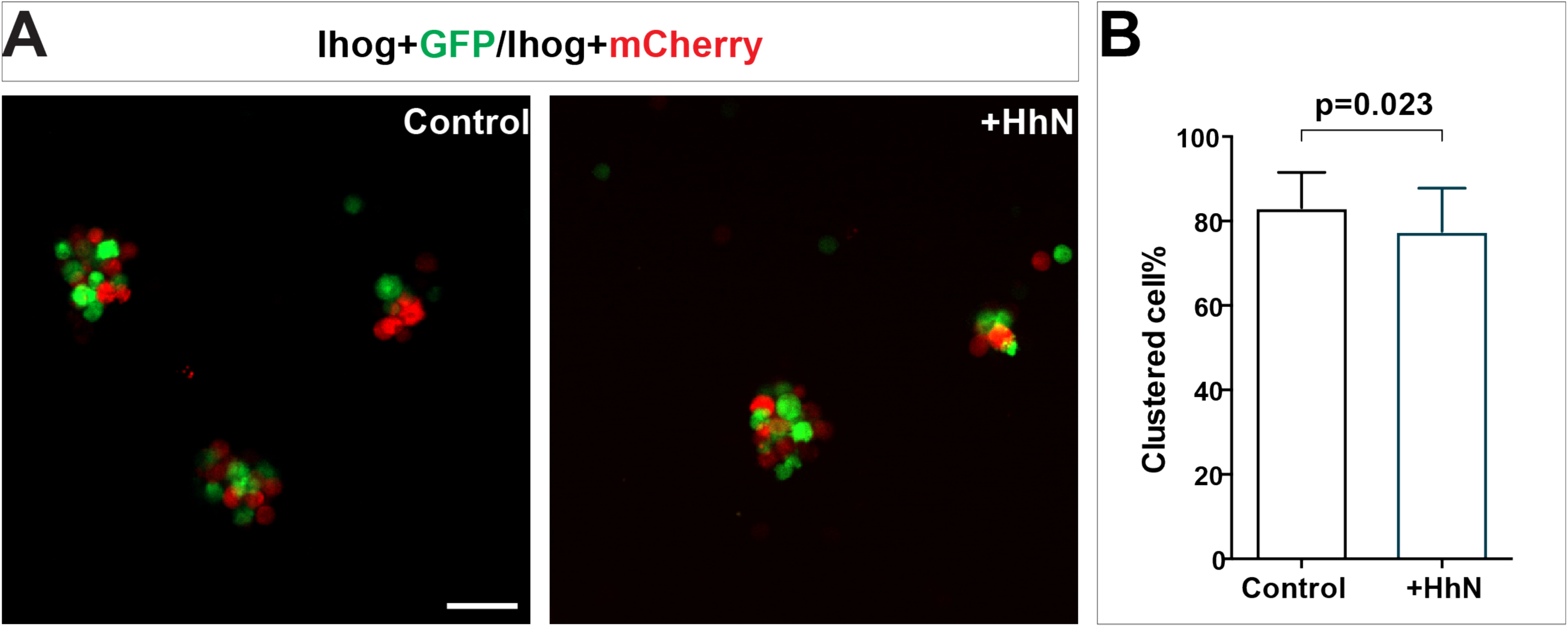
Ihog-mediated homophilic *trans* interaction in S2 cells occurs in the presence of recombinant HhN. (A) S2 cells were transfected with plasmids expressing wild-type Ihog along with GFP or mCherry. Cells were dissociated by trypsin treatment and then mixed in the absence or presence of HhN (30 μM) for 4 hr. Scale bar, 50 µm. (B) The aggregation effect from experiments like those shown in (A) was quantified as the ratio of transfected cells within a cluster to total transfected cells. Each bar shows the mean ± SD from n=30 different images. The unpaired two-tailed t-test was used for statistical analysis.

**Supplementary Figure 10.**
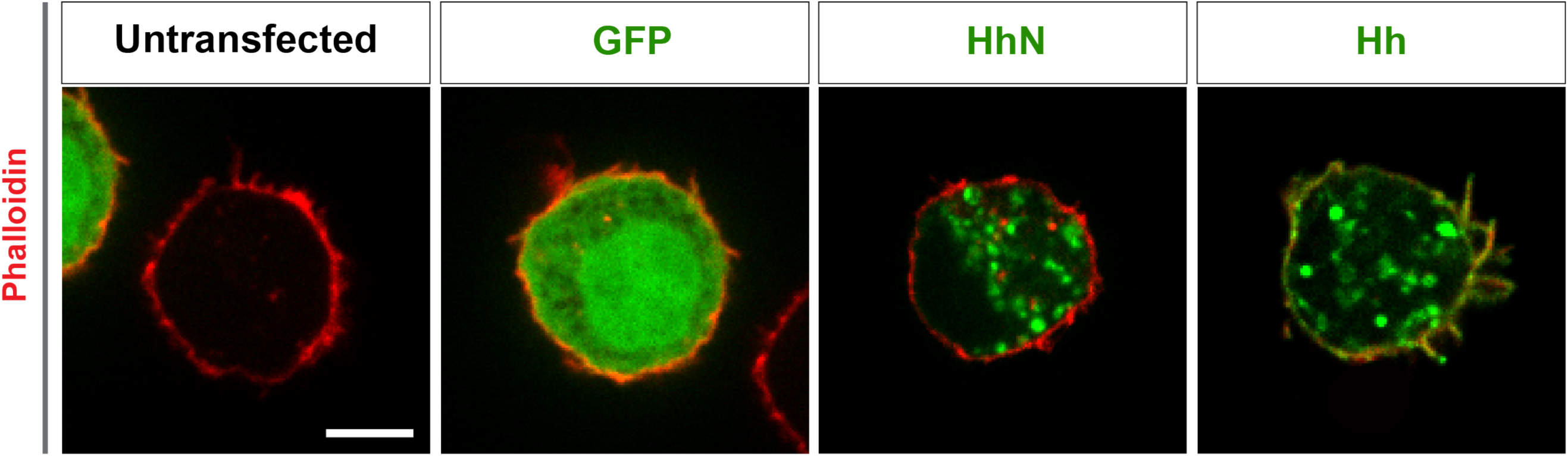
Subcellular localization of Hh and HhN in S2 cells. Untransfected S2 cell or S2 cells were transfected with constructs expressing GFP, cDNA encoding full-length Hh to generate Hh proteins that are fully processed and dual lipid modified, or cDNA encoding the amino-terminal signaling fragment HhN to generate cholesterol-free HhN. 48 hours after transfection, the S2 cells were fixed and stained with Phalloidin to detect actin (red) and the antibodies against the indicated proteins (green). Phalloidin stains filopodia-like structures and the submembrane actin network. Note that the untransfected cell is from the control cells transfected with only GFP and is from the same experiment and image. The GFP cell is partially visible in the untransfected cell image (left corner) and the nontransfected cell is partially visible in the lower right of the GFP cell image. Scale bar, 5 µm.

**Supplementary Figure 11.**
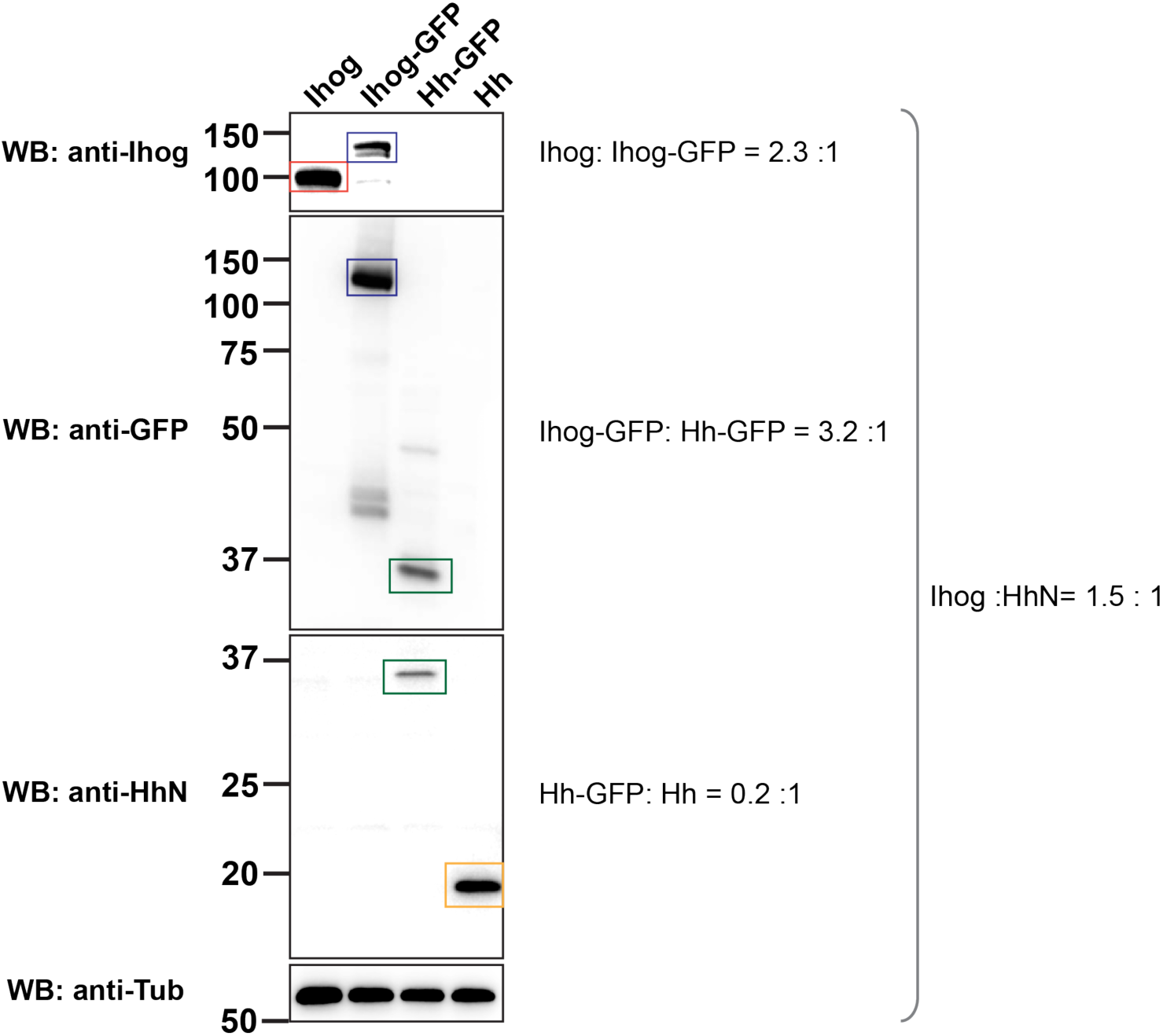
Comparison of the amount of Ihog and Hh in S2 cells analyzed in the cell-mixing assay. Aliquots of the same S2 cells analyzed in the cell mixing assay in Fig. 6 were lysed before cell mixing (Lane 1 and Lane 4). To indirectly compare the relative amounts of Ihog and Hh, lysates were collected from S2 cells that were transfected in parallel with plasmids expressing Ihog-GFP or Hh-GFP (Lane 2 and Lane 3). The amount of Ihog (red rectangle) or Hh (orange rectangle) was detected by immunoblotting with anti-Ihog or anti-Hh, respectively. The amount of GFP-tagged Ihog (blue rectangles) or GFP-tagged Hh (green rectangles) was also detected by anti-GFP. The intensity of the indicated bands was measured by ImageJ. Comparison of the intensity shows that S2 transfected cells from the same experiment used for the aggregation assays express comparable amounts (∼ 1.5:1) of Ihog and Hh proteins.

**Supplementary table 1.** Genotype of larvae for generating mosaic clones.

|  |  |
| --- | --- |
| Figure 1A | • *w; ptc-GAL4, tub-GAL80<sup>ts</sup>/UAS-GMA-GFP*<br>• *w; ptc-GAL4, tub-GAL80<sup>ts</sup>/+; UAS-Ihog/+*<br>• *w; ptc-GAL4, tub-GAL80<sup>ts</sup>/+; UAS-IhogΔFn1/+*<br>• *w; ptc-GAL4, tub-GAL80<sup>ts</sup>/+; UAS-IhogΔFn2/+*<br>• *w; ptc-GAL4, tub-GAL80<sup>ts</sup>/+; UAS-Ihog<sup>xHep</sup>/+* |
| Figure 2 | (A) *w, hs-FLP; hs-FLP/+; actin>y+>GAL4/+; UAS-Ihog/+*<br>(C) *w, hs-FLP; hs-FLP/+; actin>y+>GAL4/+; UAS-IhogΔFn2/+* |
| Figure 3A | *w, hs-FLP; hs-FLP/+; actin>y+>GAL4/+; UAS-Ihog/+*<br>*w, hs-FLP; hs-FLP/+; actin>y+>GAL4/+; UAS-IhogΔFn1/+*<br>*w, hs-FLP; hs-FLP/+; actin>y+>GAL4/+; UAS-IhogΔFn2/+*<br>*w, hs-FLP; hs-FLP/+; actin>y+>GAL4/+; UAS-Ihog<sup>xHep</sup>/+* |
| Figure 3B | *w, hs-FLP; hs-FLP/+; actin>y+>GAL4/+; UAS-Ihog/+* |
| Figure 4 | (A) *w, hs-FLP; hs-FLP/+; actin>y+>GAL4, UAS-GFP/+; UAS-Ihog/+*<br>(B-D) *w, hs-FLP/+; P{y[+t7.7] w[+mC]=CoinFLP-LexA::GAD.GAL4}attP40/UAS-IhogΔFn2-HA; P{w[+mC]=UAS-CD4-spGFP1-10}3, P{w[+mC]=lexAop-CD4-spGFP11}3/LexAop-Ihog-RFP* |
| Figure 5 | *(G) w; ptc-GAL4, tub-GAL80<sup>ts</sup>/+; UAS-Ihog/UAS-GPI-YFP* |
| Supplementary<br>Figure 1 | • *w; ptc-GAL4, tub-GAL80<sup>ts</sup>/+; UAS-myr-mRFP/+*<br>• *w; ptc-GAL4, tub-GAL80<sup>ts</sup>/+; UAS-myr-mRFP/UAS-Ihog*<br>• *w; ptc-GAL4, tub-GAL80<sup>ts</sup>/+; UAS-myr-mRFP/UAS-Ptc*<br>• *w; ptc-GAL4, tub-GAL80<sup>ts</sup>/+; UAS-myr-mRFP/UAS-Hh*<br>• *w, UAS-Dally/+; ptc-GAL4, tub-GAL80<sup>ts</sup>/+; UAS-myr-mRFP/+*<br>• *w; ptc-GAL4, tub-GAL80<sup>ts</sup>/+; UAS-myr-mRFP/UAS-Dlp* |
| Supplementary<br>Figure 2 | (A) *w; ptc-GAL4, tub-GAL80<sup>ts</sup>/+; Dally-YFP/+*<br>(B) *w; ptc-GAL4, tub-GAL80<sup>ts</sup>/+; Dally-YFP/UAS-Ihog-RFP* |
| Supplementary<br>Figure 7 | • *w; ptc-GAL4, tub-GAL80<sup>ts</sup>/+; UAS-GPI-YFP/+*<br>• *w; ptc-GAL4, tub-GAL80<sup>ts</sup>/+; UAS-lhogΔFn1/+*<br>• *w; ptc-GAL4, tub-GAL80<sup>ts</sup>/+; UAS-lhog<sup>xHep</sup>/+* |
| Supplementary<br>Figure 8 | (A) *w, boi/Y; ptc-GAL4, tub-GAL80<sup>ts</sup>/+; UAS-myr-mRFP/Tm6B*<br>(B) *w, boi/Y; ptc-GAL4, tub-GAL80<sup>ts</sup>/+; UAS-myr-mRFP/UAS-lhog-RNAi* |

